# Discovery of an embryonically derived bipotent population of endothelial-macrophage progenitor cells in postnatal aorta

**DOI:** 10.1101/2021.10.18.464001

**Authors:** Anna E. Williamson, Sanuri Liyanage, Mohammadhossein Hassanshahi, Malathi S.I. Dona, Deborah Toledo-Flores, Dang X.A. Tran, Catherine Dimasi, Nisha Schwarz, Sanuja Fernando, Thalia Salagaras, Aaron Long, Jan Kazenwadel, Natasha L. Harvey, Grant R. Drummond, Antony Vinh, Vashe Chandrakanthan, Ashish Misra, Zoltan Neufeld, Joanne T.M. Tan, Luciano Martelotto, Jose M. Polo, Claudine S. Bonder, Alexander R. Pinto, Shiwani Sharma, Stephen J. Nicholls, Christina A. Bursill, Peter J. Psaltis

## Abstract

Converging evidence indicates that extra-embryonic yolk sac is the source of both macrophages and endothelial cells in adult mouse tissues. Prevailing views are that these embryonically derived cells are maintained after birth by proliferative self-renewal in their differentiated states. Here we identify clonogenic endothelial-macrophage (EndoMac) progenitor cells in the adventitia of embryonic and postnatal mouse aorta, that are independent of Flt3-mediated bone marrow hematopoiesis and derive from an early embryonic CX_3_CR1^+^ and CSF1R^+^ source. These bipotent progenitors are proliferative and vasculogenic, contributing to adventitial neovascularization and forming perfused blood vessels after transfer into ischemic tissue. We establish a regulatory role for angiotensin II, which enhances their clonogenic and differentiation properties and rapidly stimulates their proliferative expansion *in vivo*. Our findings demonstrate that embryonically derived EndoMac progenitors participate in local vasculogenic responses in the aortic wall by contributing to the expansion of endothelial cells and macrophages postnatally.

## INTRODUCTION

Among diverse roles, macrophages are integral to development of blood and lymphatic vessels during normal organogenesis and responses to tissue injury, ischemia and other diseases ^1–4^. They proliferate and assemble around neovessels, producing angiogenic factors and supporting endothelial anastomoses and vascular remodeling ^2^. In return, endothelial cells help regulate the self-renewal of hematopoietic stem cells (HSCs) and their differentiation into macrophages ^5^. Understanding how macrophage-endothelial interactions arise is important to target inflammation and neovascularization in different pathophysiological conditions.

Historically, circulating monocytes were thought to be the source of tissue macrophages ^6^. Monocytes derive from definitive hematopoiesis, which originates embryonically with emergence of HSC clusters from the endothelium of the aorta-gonad-mesonephros (AGM) at around embryonic day (E) 10.5 in mice ^7–9^. These HSCs seed fetal liver before colonizing bone marrow (BM) perinatally, the main hematopoietic organ after birth. Numerous studies have now established that HSC-monocyte ancestry does not account for all tissue macrophages ^10–16^. During embryogenesis, extra-embryonic yolk sac (YS) is the first site to produce macrophages via distinct developmental programs. This begins with primitive macrophage precursors in YS blood islands between E7.0 and E8.25 ^17, 18^, followed by erythromyeloid progenitors (EMPs) which bud from specialized YS hemogenic endothelium ^19^. From E7.5, c-Kit^+^CSF1R^+^ EMPs produce erythrocytes, megakaryocytes and YS macrophages in a process that is independent of the transcription factor, Myb ^11, 16, 20^. This does not involve monocyte intermediates but rather sequential differentiation into CX_3_CR1^+^ pre-macrophages and mature F4/80^Hi^ macrophages ^17, 21–23^. Macrophage progenitors, in both multipotent EMP and pre-macrophage stages, expand in YS and circulate to embryonic tissues to complete their maturation ^24^. This migration peaks around E10.5 and is mostly complete by E12.5. From E8.5, Myb^+^c-Kit^+^CSF1R^Lo^ EMPs also differentiate into YS macrophages and traffic to liver, where they expand and generate lineage-specific hematopoietic progenitors and fetal blood cells, including monocytes ^23, 25^. Long-lived populations of embryonically derived macrophages persist in adult tissues, including brain, skin, liver, heart, lung and aortic adventitia ^10–16, 26^. These are maintained independently of BM hematopoiesis through local proliferation and seemingly, by self-renewal ^27^.

Recent studies indicate that YS EMPs also contribute endothelial cells to the blood and lymphatic vessels of some organs ^4, 28^. Although disputed ^29^, the possibility that EMPs produce both hematopoietic and endothelial cells is intriguing, especially given long-standing speculation around the embryonic existence of mesoderm-derived hemangioblasts ^30^. While hemangioblasts have been shown to emerge during hemato-endothelial differentiation of pluripotent stem cells *in vitro* ^31, 32^, their presence in postnatal tissues remains unproven ^33^.

Here, we identify clonogenic endothelial-macrophage (EndoMac) progenitor cells in postnatal aorta, that are embryonically derived from YS and are seeded around E10.5 of gestation. These highly proliferative progenitors are independent of Flt3-mediated hematopoiesis and are a local source of macrophage and endothelial renewal and expansion during adventitial neovascularization.

## RESULTS

### c-Kit and CX_3_CR1 identify CFU-M progenitors in postnatal aorta

Previously, we identified that the adventitia of mouse arteries contains cells with hematopoietic potential that selectively produce macrophage colony-forming units (CFU-M) in methylcellulose ^34, 35^. To establish the true nature of these CFU-M forming cells, we began by culturing aortic digests from 12-week-old (w) C57BL/6J mice in methylcellulose (MethoCult GF M3434, StemCell Technologies) for 14 days (d). This generated 17.0±12.0 CFU-M/10^5^ cells (n=10), with colonies further classified based on size as ~77% small (~30-100 cells), ~23% medium (~100-1000 cells) and <1% large (>1000 cells) (**Fig. 1A**). Data from other studies have been interpreted to support the self-renewal of mature macrophages in postnatal aorta ^15, 36^. We therefore investigated whether adult aortic CFU-M can renew *in vitro*. CFU-M were isolated and disaggregated, with their content replated in single-cell secondary (2°) cultures for another 14 d in methylcellulose. Daily inspection revealed that single cells from small CFU-M gave rise to 2° clusters (<30 cells), whereas those from medium CFU-M generated small 2° colonies (**Fig. 1B**), with renewal frequency of 1 new colony produced per ~15 cells (mean 14.8±16.3, n=20 replicates; also see Source Data for **Extended Data Fig. 4L**). We observed no further colony formation in tertiary cultures, indicating that the renewal ability of CFU-M from adult C57BL/6J aortas is finite.

**Fig. 1.**
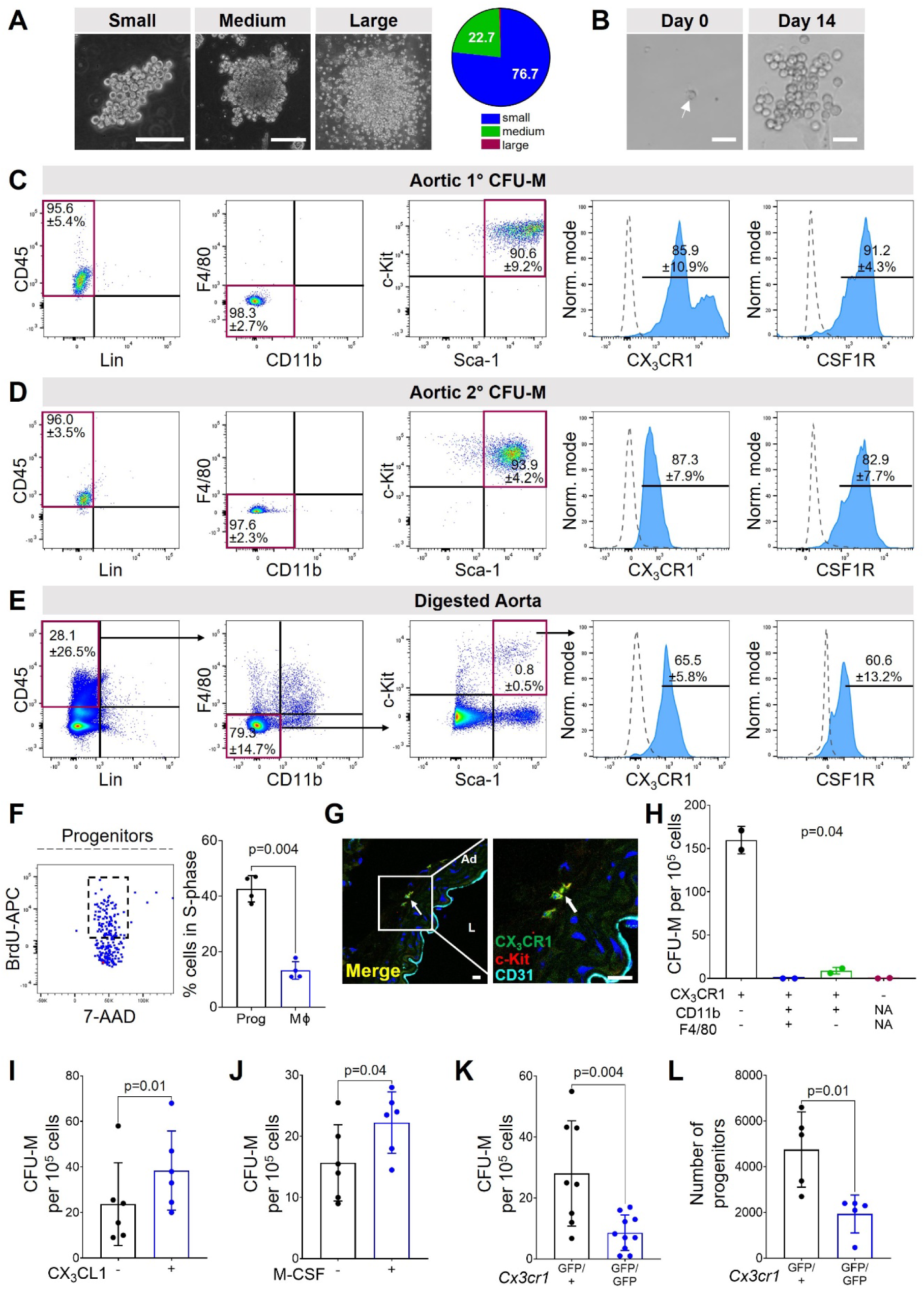
Immunophenotypic characterization of CFU-M progenitors in adult aorta. (A) Examples of small, medium and large CFU-M from 12 w C57BL/6J aortic cells. Pie chart shows breakdown of CFU-M by colony size (n=5). (B) Representative example of renewal of 2° CFU-M from a single cell plated from a medium 1° CFU-M. (C, D) Flow cytometry of cells from (C) 1° and (D) 2° aortic CFU-M of medium size (n≥3 for each marker). Blue histogram, sample. Dotted histogram, Fluorescence-minus-one (FMO) control. (E) Flow cytometry shows presence of progenitors in 12 w C57BL/6J aorta *in vivo* (n=6 for each marker with 1-2 mice per replication). (F) Flow cytometry of bromodeoxyuridine (BrdU) uptake versus 7-aminoactinomycin (7-AAD) labeled cells gated from aortic progenitors *in vivo*. Dotted box, S-phase. Graph shows percentages of progenitors (Prog) and macrophages (Mϕ) in S-phase from 12 w C57BL/6J aorta 24 h after administration of BrdU (n=4; paired t-test). (G) Confocal microscopy of immunolabeled section of descending aorta from 12 w *Cx3cr1*^GFP/+^ mouse shows adventitial (Ad) CX_3_CR1^+^c-Kit^+^ progenitors. L, lumen. (H) CFU-M yield from FACS-isolated cells from adult *Cx3cr1*^GFP/+^ aortas (two experiments, n=6 mice each). N/A, not applicable. (I, J) CFU-M yield from 12 w C57BL/6J aortic cells in presence of (I) 100 nM CX_3_CL1 or (J) 50 nM M-CSF (n=6; paired t-tests). (K, L) Frequency of (K) CFU-M (n=8-10) and (L) progenitors (n=5) from aortas of 12 w *Cx3cr1*^GFP/+^ and *Cx3cr1*^GFP/GFP^ mice. Unpaired t-test for (K) and Mann-Whitney test for (L). Data summarized as mean±SD. Scale bar, 100 µm in (A), (B) and 20 µm in (G). Also see **Extended Data Fig. 1–3**.

Having established that medium CFU-M form 2° colonies from a single-cell source, we used flow cytometry to examine their composition. This revealed a population in both 1° and 2° CFU-M that was predominantly negative for lineage (Lin) and monocyte/macrophage markers (CD11b, F4/80), but positive for CD45, Sca-1, c-Kit, CX_3_CR1 and CSF1R (**Fig. 1C, D** and **Extended Data Fig. 1A**). In contrast, large CFU-M which were only rarely produced from adult aorta, contained 32.0±8.8% CD45^+^CD11b^+^F4/80^+^ macrophages (n=6). These were located on the outer perimeter of large colonies, while the inner core was made up of Lin^−^ CD45^+^CD11b^−^F4/80^−^Sca-1^+^c-Kit^+^ progenitors. Medium CFU-M also expressed the hematopoietic progenitor and EMP markers, CD34, CD16/32, CD93 (AA4.1) and CD43 ^37^, but not CD41 (very early-stage hematopoiesis) or Flt3 (CD135) (**Extended Data Fig. 1B**).

The stem cell marker, c-Myc, which regulates self-renewal was expressed in 15.0±5.4% of permeabilized cells, while Sox2, Nanog and Oct4 which are also associated with pluripotency, were very lowly expressed. Other mature myeloid markers were negative (<1%) (CD64, CD86, MerTK) or lowly expressed (CD24, MHC II, CCR2, Ly6C, Gr1, LYVE-1). Medium CFU-M were also negative for the erythroid marker, TER-119, and endothelial marker, CDH5 (CD144), with <5% surface expression of VEGFR2 and TIE2; however, they did express CD31, which is also found on myeloid progenitors ^17^ (**Extended Data Fig. 1B**). Immunofluorescence labeling and confocal microscopy also showed that progenitors in aortic CFU-M were proliferating (Ki67^+^) and homogeneously c-Kit^+^, but were negative for the macrophage marker, CD68, and endothelial marker, Endomucin (EMCN) (**Extended Data Fig. 2A**).

Importantly, we identified Lin^−^CD45^+/Lo^CD11b^−^F4/80^−^Sca-1^+^c-Kit^+^ progenitors in 12 w C57BL/6J aorta *in vivo* (0.15±0.13% or 1,632±495 cells per aorta, n=6), resembling the surface phenotype of CFU-M (**Fig. 1E** and **Extended Data Fig. 1A**). Bromodeoxyuridine (BrdU) uptake 24 h after administration revealed that ~43% of these progenitors were in S-phase of cell cycle, compared to ~13% of macrophages in donor-matched aortas (**Fig. 1F**). We also used fluorescence activated cell sorting (FACS) of aortic digests from C57BL/6J mice to isolate CD45^+^CD11b^−^F4/80^−^Sca-1^+^c-Kit^+^ progenitors and confirmed their ability to form CFU-M (**Extended Data Fig. 2B-D**). These colonies had similar surface marker expression as the medium CFU-M grown from unfractionated aortic cells (**Extended Data Fig. 2E**). In contrast, no colonies grew from isolated CD45^+^CD11b^+^F4/80^+^ macrophages.

Fractalkine receptor, CX_3_CR1, has been used to characterize tissue-resident macrophages, including those in adventitia, and to trace their embryonic origins ^12, 14, 15^. As CFU-M progenitors expressed CX_3_CR1, we studied *Cx3cr1*^GFP/+^ mice which express green fluorescent protein (GFP) under control of the *Cx3cr1* locus ^38^. CFU-M from *Cx3cr1*^GFP/+^ aortas were GFP^+^ under fluorescence microscopy and flow cytometry (**Extended Data Fig. 3A**). Whereas other studies have focused on CX_3_CR1 expression by aortic macrophages ^15^, we also identified CX_3_CR1^+^c-Kit^+^ progenitors in aortic adventitia (**Fig. 1G**) and determined by flow cytometry that they accounted for 18.9±4.7% of CX_3_CR1^+^ cells in the aortic wall (n=6) (**Extended Data Fig. 3B,C**). FACS isolation of aortic cells from *Cx3cr1*^GFP/+^ mice showed that CFU-M forming capacity was restricted to GFP^+^ (especially GFP^Bright^) cells (**Extended Data Fig. 3D**), and specifically GFP^+^CD11b^−^F4/80^−^ progenitors, not GFP^+^ macrophages (**Fig. 1H** and **Extended Data Fig. 3E,F**).

As evidence of the functional importance of CX_3_CR1, the addition of its ligand, CX_3_CL1, resulted in 1.6-fold higher CFU-M yield from aortic cells (**Fig. 1I**). This was similar to the effect of macrophage colony-stimulating factor (M-CSF), the ligand for CSF1R (**Fig. 1J**). We also compared *Cx3cr1*^GFP/+^ mice to *Cx3cr1*^GFP/GFP^ littermates, that lack both functional *Cx3cr1* alleles. *Cx3cr1*^GFP/GFP^ aortas had lower CFU-M yield (**Fig. 1K**) and contained ~60% fewer progenitors by flow cytometry (**Fig. 1L**). Therefore CX_3_CR1 is both a marker of CFU-M progenitors and promotes their clonogenicity and prevalence in aorta.

### Aortic CFU-M progenitors are independent of Flt3^+^ BM hematopoietic progenitors

C-C chemokine receptor 2 (CCR2), the cognate receptor for C-C chemokine ligand 2 (CCL2), has been used to differentiate between monocyte-derived (CCR2^+^) and locally maintained, embryonically derived (CCR2^−^) macrophages in some tissues ^39^. Unlike CX_3_CR1, CCR2 was lowly expressed on aortic CFU-M progenitors (**Extended Data Fig. 4A**). Aortic cells from *Ccr2*^−/−^ mice formed more CFU-M than wildtype littermates, with a higher percentage of progenitors (**Extended Data Fig. 4B-D**). Similarly, when clodronate liposomes were used to deplete blood monocytes, we observed a significant increase in aortic CFU-M formation (**Extended Data Fig. 4E-G**). Both these results suggest that the size of the aortic progenitor population is not dependent on circulating monocytes.

This led us to examine the relationship between CFU-M progenitors and BM hematopoiesis by using *Flt3*^Cre^ x *Rosa*^mT/mG^ mice. BM hematopoietic progenitors transiently upregulate the receptor tyrosine kinase Flt3 during hematopoietic differentiation ^40^. In *Flt3*^Cre^ x *Rosa*^mT/mG^ mice cells that originate from BM hematopoietic progenitors express GFP (Flt3-Cre^+^), while those that do not are GFP^−^ (Flt3*-*Cre^−^) ^41^. We observed high Flt3-Cre labeling efficiency of CD45^+^ Flt3/CD135^+^ cells in adult BM in these mice with 85.9±2.7% GFP^+^ (**Extended Data Fig. 4H**). Consistently, whereas BM produced a mixture of CFU-M and non-macrophage colonies that were ~90% Flt3-Cre^+^, adult aortic CFU-M from *Flt3*^Cre^ x *Rosa*^mT/mG^ mice were ~95% Flt3-Cre^−^ (**Fig. 2A,B**). Similarly, >90% of Lin^−^CD45^+/Lo^CD11b^−^F4/80^−^Sca-1^+^c-Kit^+^ progenitors in aorta were Flt3-Cre^−^ (**Fig. 2C**). In contrast, **~**65% of short-term HSCs (ST-HSCs) and ~95% of multipotent progenitors (MPPs) in BM, and ~95% of monocytes in peripheral blood were Flt3-Cre^+^, as were ~40% of long-term BM HSCs (LT-HSCs) (**Fig. 2D** and **Extended Data Fig. 4I,J**). In keeping with previous studies ^15, 36^, aorta contained a mixture of Flt3-Cre^+^ (~85%) and Flt3-Cre^−^ (~15%) macrophages (**Extended Data Fig. 4K**). For reference, macrophages in BM and spleen were >95% Flt3-Cre^+^ and brain microglia ~94% Flt3-Cre^−^. Therefore, aortic CFU-M progenitors are unlikely to be derived from Flt3-mediated BM hematopoiesis.

**Fig. 2.**
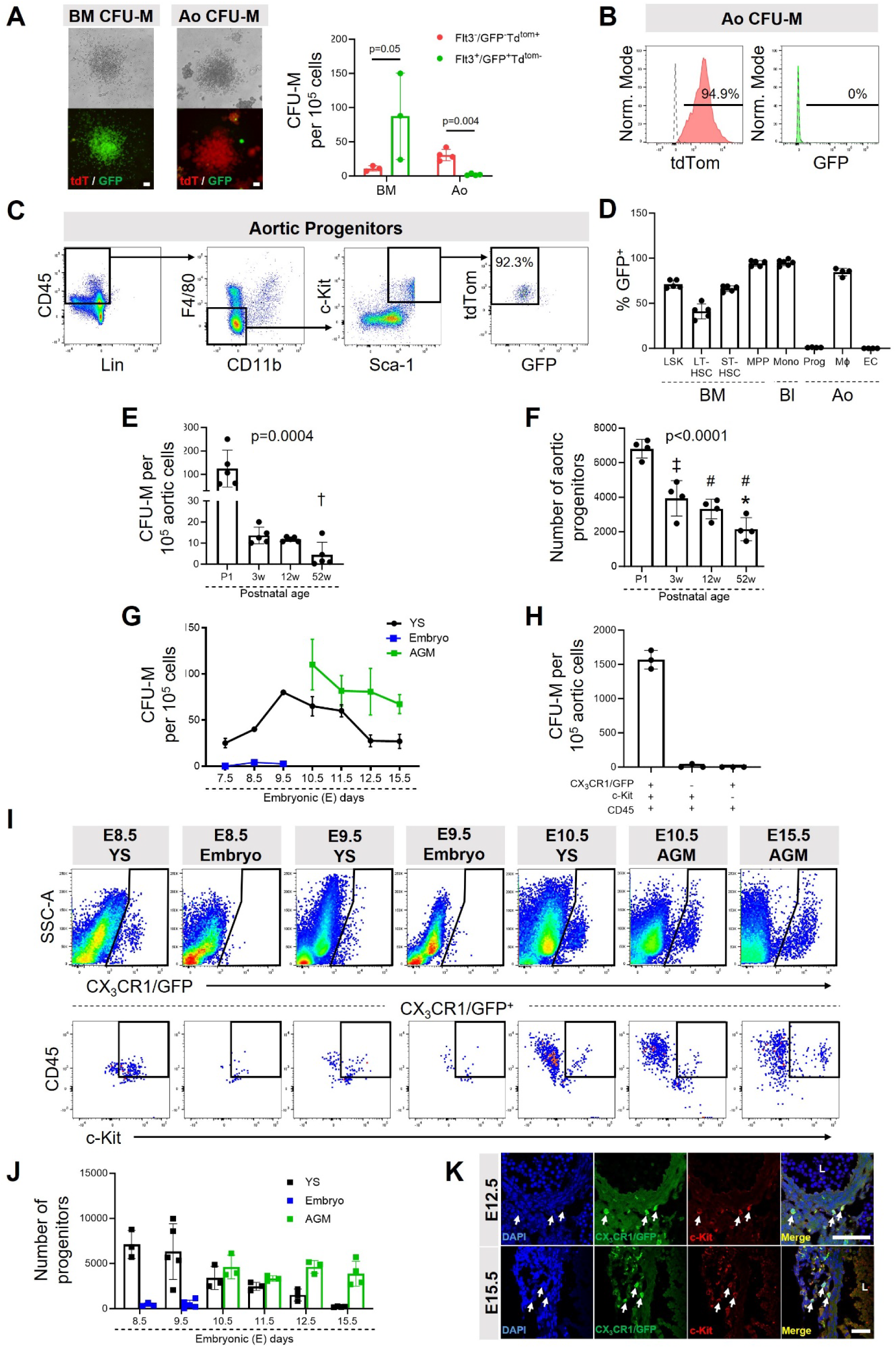
Aortic CFU-M progenitors are independent of Flt3-mediated hematopoiesis and are seeded embryonically. (A) Light and fluorescence microscopy images of CFU-M from BM and aorta (Ao) of adult *Flt3*^Cre^ x *Rosa*^mT/mG^ mice. Graph shows Flt3^+^ (green) and Flt3^−^ (red) CFU-M yield from BM and Ao (n=3-4; paired t-test for each tissue). (B) Flow cytometry histograms showing tdTom and GFP expression of cells in CFU-M from *Flt3*^Cre^ x *Rosa*^mT/mG^ aorta (n=1). Dotted histogram, C57BL/6J control. (C) Flow cytometry plots show GFP^−^tdTom^+^ (Flt3-cre^−^) status of aortic progenitors from adult *Flt3*^Cre^ x *Rosa*^mT/mG^ mice. Percentage represents mean of n=4. *Please see Source Data file for FMO controls.* (D) Comparison of proportion of GFP^+^ (Flt3-cre^+^) cells in BM long-term (LT) and short-term (ST) HSCs, multipotent progenitors (MPPs), blood monocytes (Mono) and aortic progenitors (Prog), macrophages (Mϕ) and endothelial cells (ECs) from adult *Flt3*^Cre^ x *Rosa*^mT/mG^ mice (n=4-6). LSK, Lin^−^Sca-1^+^c-Kit^+^; Bl, blood. *Please see Methods for immunophenotypic definitions of BM cell populations*. (E, F) Comparisons of (E) CFU-M yield (n=5; Kruskal-Wallis test) and (F) number of progenitors (n=4; one-way ANOVA) from C57BL/6J aortas at different ages. Multiple comparison tests: *p<0.05 for 3 w vs 52 w; ‡p<0.001 and #p<0.0001 for comparison to P1. (G) CFU-M yield from different embryonic tissues of *Cx3cr1*^GFP/+^ mice at different embryonic ages (n=1-12). (H) CFU-M yield from different subpopulations of FACS-isolated cells from E9.5 YS from *Cx3cr1*^GFP/+^ mice (n=3 experiments, each using ≥6 pooled YS). Repeated measures ANOVA p=0.003. (I) Flow cytometry of digests of YS, whole embryo and AGM from *Cx3cr1*^GFP/+^ mice at different embryonic ages showing CX_3_CR1/GFP^+^ cells (top row) and CX_3_CR1/GFP^+^c-Kit^+^CD45^+^ progenitors (bottom row) (n=3-5). (J) Number of CX_3_CR1/GFP^+^c-Kit^+^CD45^+^ progenitors in YS, whole embryo or AGM at different embryonic ages (n=3-5). (K) Confocal microscopy of immunolabeled E12.5 and E15.5 AGM from *Cx3cr1*^GFP/+^ mice shows adventitial CX_3_CR1/GFP^+^c-Kit^+^ progenitors (arrows). L, lumen. Data summarized as mean or mean±SD. Scale bar, 100 μm in (A) and 50 μm in (K). Also see **Extended Data Fig. 4** and **5**.

### Aortic CFU-M progenitors are seeded embryonically

To further elucidate the origins of aortic CFU-M progenitors, we performed age profiling after birth. Aortic CFU-M yield was ~10-fold higher from P1 (postnatal day 1) than 3 w and 12 w mice, and lowest from 52 w mice (**Fig. 2E**). P1 aortas produced more large and medium colonies, with the latter showing greater ability to form 2° colonies after single-cell replating than was the case from older mice (**Extended Data Fig. 4L,M**). Flow cytometry supported the higher abundance of progenitors in P1 aorta and their declining prevalence as mice age (**Fig. 2F**).

As aortic CFU-M progenitors are present at birth, we examined their emergence during embryonic development. CFU-M grew from digests of YS from E7.5, with their yield peaking at E9.5 (**Fig. 2G** and **Extended Data Fig. 4N**). 2° renewal capacity was 100% with one new colony formed for each cell plated from medium CFU-M from E9.5 YS (**Extended Data Fig. 4M**). Although CFU-M were rare from digests of whole embryo between E7.5 and E9.5, they were more prevalent from AGM from E10.5 onwards (**Fig. 2G**). Coinciding with the emergence of definitive HSCs from dorsal aorta at ~E10.5-11.5 ^9^, we also observed non-macrophage colonies (G, granulocyte; GM, granulocyte-macrophage; GEMM, granulocyte-erythrocyte-monocyte-megakaryocyte; BFU-E, burst-forming units-erythroid) from AGM during this gestational window (**Extended Data Fig. 4O**). However, unlike CFU-M these disappeared by E12.5. Colony growth from liver increased between E10.5 and E11.5 and consisted mostly of non-macrophage colonies through to E15.5, consistent with seeding of HSCs and multipotent progenitors from AGM and YS ^23, 25^ (**Extended Data Fig. 4P**).

We next investigated which cells in early YS are responsible for producing CFU-M. Using *Cx3cr1*^GFP/+^ embryos, three populations were FACS isolated from E9.5 YS: 1) CX_3_CR1/GFP^+^c-Kit^+^CD45^+^, 2) CX_3_CR1/GFP^−^c-Kit^+^CD45^+^ and 3) CX_3_CR1/GFP^+^c-Kit^−^ CD45^+^ cells (**Extended Data Fig. 5A**). CFU-M were selectively produced by the CX_3_CR1/GFP^+^c-Kit^+^CD45^+^ fraction (**Fig. 2H** and **Extended Data Fig. 5B**). These CFU-M expressed surface markers consistent with those of CFU-M from adult aorta (**Extended Data Fig. 5C**) and were able to renew from a single-cell source with 78% (7 out of 9 replicate wells) and 14% (1 out of 7) efficiency after 2° and 3° replating, respectively. In contrast, CX_3_CR1/GFP^−^c-Kit^+^CD45^+^ cells produced mostly non-macrophage colonies, while no colonies grew from CX_3_CR1/GFP^+^c-Kit^−^CD45^+^ cells (**Fig. 2H** and **Extended Data Fig. 5B**). We also observed the presence of CX_3_CR1/GFP^+^c-Kit^+^CD45^+^ progenitors in YS from E8.5 and AGM from E10.5, in keeping with CFU-M growth (**Fig. 2I,J** and **Extended Data Fig. 5D,E**). Lastly, using confocal microscopy, we identified CX_3_CR1/GFP^+^c-Kit^+^ progenitors in aortic adventitia at E12.5 and E15.5 (i.e., after disappearance of definitive HSCs from AGM) (**Fig. 2K**).

We then used *Flt3*^Cre^ x *Rosa*^mT/mG^ mice again to determine whether CFU-M producing progenitors in YS arise from a Flt3^+^ source. As for adult BM, we observed high Flt3-Cre labeling efficiency in E10.5 YS, the site where Flt3 surface expression emerges ^42^, with 81.4±3.9% of CD45^+^Flt3/CD135^+^ cells expressing GFP (n=5) (**Extended Data Fig. 5F**). Focusing on c-Kit^+^CD45^+^ progenitors, most of these cells were Flt3/CD135^−^ and GFP^−^ (88.9±3.5%), whereas only a small proportion were Flt3/CD135^+^ and GFP^+^ (6.5±3.3%) (**Extended Data Fig. 5G**). Additionally, we sorted c-Kit^+^CD45^+^Flt3/CD135^−^ and c-Kit^+^CD45^+^Flt3/CD135^+^ cells from E10.5 YS from C57BL/6J embryos and performed the CFU assay. Notably, CFU-M predominantly grew from c-Kit^+^CD45^+^Flt3/CD135^−^ cells (**Extended Data Fig. 5H,I**). Therefore, at least the majority of CFU-M producing progenitors in YS arise from a Flt3^−^ source, consistent with the independence of aortic progenitors from Flt3-mediated BM hematopoiesis, described above.

### Aortic CFU-M progenitors originate from CX_3_CR1^+^ and CSF1R^+^ embryonic progenitors

Murine aortic adventitia has been shown to contain locally maintained macrophages derived from CX_3_CR1^+^ and CSF1R^+^ YS progenitors ^15^. As CFU-M progenitors appear in YS before the onset of definitive hematopoiesis in E10.5 AGM, we used timed fate-mapping approaches at E8.5 and E9.5 to study their embryonic origins. Female *Cx3cr1*^CreER-YFP^ mice, which express Cre recombinase under control of the *Cx3cr1* promoter upon exposure to 4-hydroxytamoxifen (TAM), were crossed to male *Rosa*^tdTom^ mice ^15^. Pregnant dams were administered TAM at E8.5 or E9.5 to induce irreversible expression of the tdTomato (tdTom) reporter in CX_3_CR1^+^ cells and their progeny. Whereas previous studies have focused on the macrophage ^11, 15^ or endothelial fate ^28, 29^ of YS progenitors, we tracked both lineages in aorta together with CFU-M progenitors. After E8.5 induction, adult aorta contained tdTom^+^ cells that included all three populations, with macrophages being most prevalent, followed by progenitors (**Extended Data Fig. 6A,B**). For E9.5 induction, we first confirmed that there was negligible labeling of cells in BM at 12 w, the main site of definitive hematopoiesis postnatally (**Fig. 3A**). Consistently, CFU-M from adult BM were exclusively tdTom^−^, whereas tdTom^+^ CFU-M were produced by E15.5 AGM cells and 12 w aortic cells, indicating their origins from an E9.5 CX_3_CR1^+^ source (**Fig. 3B**). In keeping with this, we identified tdTom^+^ progenitors, as well as tdTom^+^ macrophages and endothelial cells in both E15.5 AGM and 12 w aorta, although tdTom^+^ progenitors were less frequent postnatally (**Fig. 3C, D**). After normalizing tdTom labeling to results for brain microglia, E9.5 CX_3_CR1^+^ cells accounted for the source of ~43% of progenitors, ~32% of macrophages and ~19% of endothelial cells in 12 w aorta, while making negligible contribution to monocytes in blood or aorta (**Fig. 3E,F**). Immunofluorescent confocal microscopy also confirmed the presence of intimal and adventitial tdTom^+^CDH5^+^ endothelial cells, and adventitial tdTom^+^CD68^+^ macrophages and tdTom^+^c-Kit^+^ progenitors in aortas of E9.5 TAM-induced adult *Cx3cr1*^CreER-YFP^ x *Rosa*^tdTom^ mice (**Fig. 3G**).

**Fig. 3.**
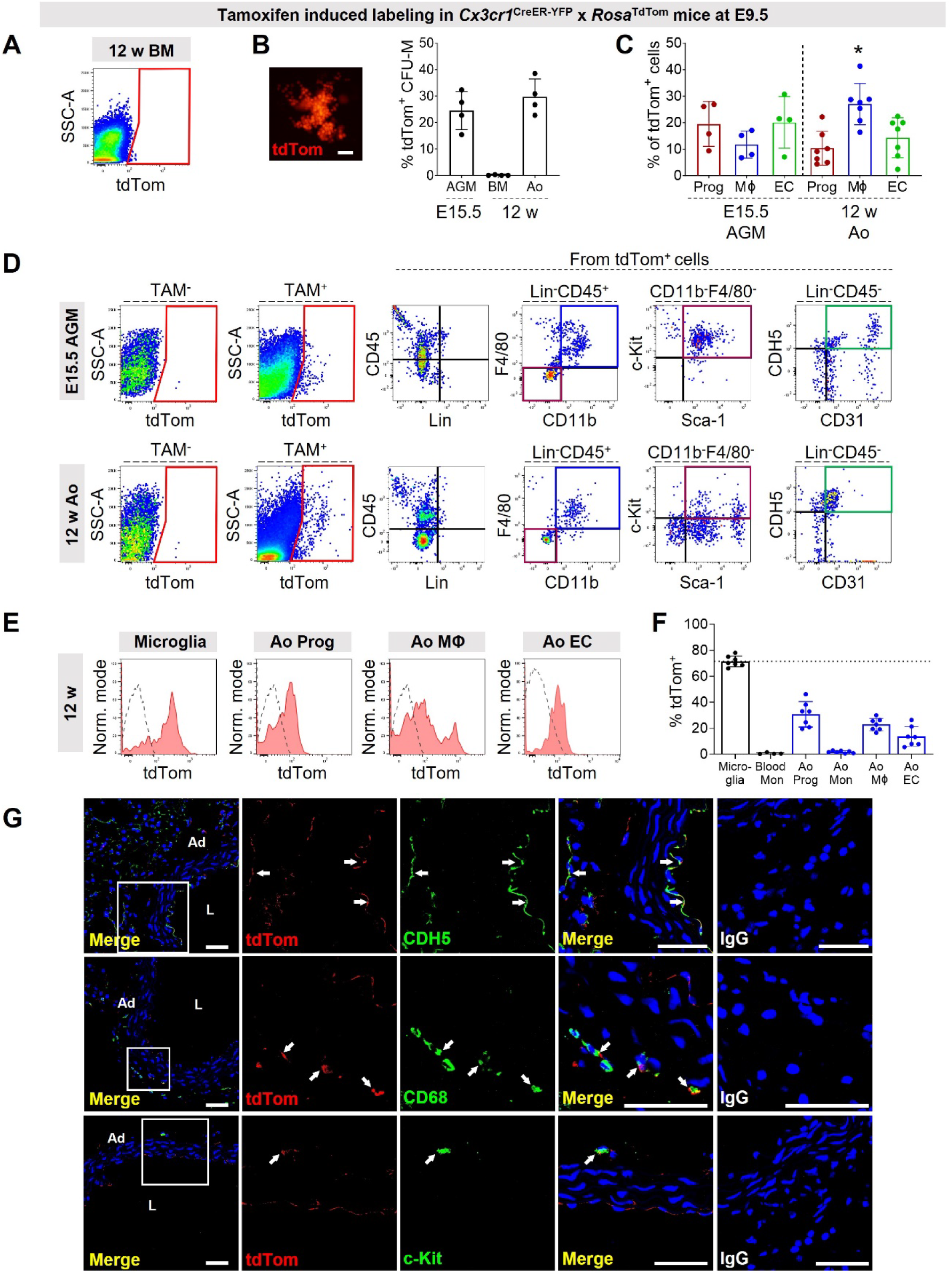
Aortic CFU-M progenitors arise from an early embryonic CX_3_CR1*^+^* source. (A-F) 4-hydroxytamoxifen (TAM)-induced labeling was performed in *Cx3cr1*^CreER-YFP^ x *Rosa*^tdTom^ mice at E9.5 for subsequent analysis. (A) Representative plot shows negligible tdTom expression in 12 w bone marrow (BM) (n=3-4). (B) Image of tdTom^+^ CFU-M from 12 w aorta. Graph shows % of tdTom^+^ CFU-M from different tissues and ages (n=4). Ao, aorta. AGM, aorta-gonad-mesonephros. (C, D) Graph and flow cytometry plots show composition of tdTom^+^ cells in E15.5 AGM and 12 w aorta, with gated regions showing progenitors (Prog, maroon), macrophages (Mφ, blue) and endothelial cells (EC, green) (n=4-7). tdTom expression from *Cx3cr1*^CreER-YFP^ x *Rosa*^tdTom^ mice which did not receive TAM is also shown as negative control. Repeated measures ANOVA p=0.004 for 12 w; Multiple comparison tests: *p<0.05 for Mφ vs Prog and EC. (E, F) Histograms and graph show % tdTom expression in different populations from 12 w mice (n=4-7). Red histogram, sample. Dotted histogram, TAM^−^ control. Mon, monocytes. (G) Confocal microscopy images of immunolabeled sections of adult aorta from E9.5 TAM-induced *Cx3cr1*^CreER-YFP^ x *Rosa*^tdTom^ mice show VE-cadherin (top row), CD68 (middle row) and c-Kit (bottom row) expression in tdTom^+^ cells (arrows). Merged images of IgG controls for each labeling are also shown. Ad, adventitia. L, lumen. *Please see Source Data file for larger version of these images.* Data summarized as mean±SD. Scale bar, 100 μm in (B) and 40 μm in (G). Also see **Extended Data Fig. 6** and **7**.

Complementary fate-mapping was performed by giving TAM to *Csf1r*^Mer-iCre-Mer^ x *Rosa*^mT/mG^ mice at E8.5 to induce GFP expression in YS CSF1R^+^ cells, including EMPs and their progeny ^14^. Approximately 30% of CFU-M from 12 w aorta were GFP^+^, compared to <2% from donor-matched BM (**Extended Data Fig. 6C**). Using flow cytometry, we identified GFP^+^ progenitors, macrophages and endothelial cells in adult aorta (**Extended Data Fig. 6D**). After normalizing to labeling of microglia, ~62% of progenitors, ~26% of macrophages and ~12% of endothelial cells in adult aorta were from an E8.5 CSF1R^+^ source, consistent with YS EMP origins (**Extended Data Fig. 6E,F**). Immunolabeling and confocal microscopy also demonstrated GFP^+^ cells in ascending and descending aorta, comprising intimal and adventitial CD31^+^ endothelium and adventitial c-Kit^+^ progenitors and CD68^+^ macrophages (**Extended Data Fig. 6G,H**).

We next used FACS to isolate embryonically derived tdTom^+^ or GFP^+^ progenitors, macrophages and endothelial cells from aortas of E9.5 TAM-induced 12 w *Cx3cr1*^CreER-YFP^ x *Rosa*^tdTom^ mice or E8.5 TAM-induced 12 w *Csf1r*^MerCreMer^ x *Rosa*^mT/mG^ mice, respectively (**Extended Data Fig. 7A,B**). Wright-Giemsa staining showed that progenitors had rounded morphology with smooth surface membrane and lacked the protrusions and intracellular vacuolations of macrophages (**Extended Data Fig. 7C**). Although they had similarly sized nuclei as macrophages, progenitors had less cytoplasm and a higher nuclear:cell area ratio (**Extended Data Fig. 7C**). CFU-M were only produced by progenitors and not macrophages or endothelial cells after culture in methylcellulose (**Extended Data Fig. 7D**). These data collectively indicate that aortic CFU-M progenitors originate from an early embryonic CSF1R^+^ and CX_3_CR1^+^ source.

### CFU-M progenitors have endothelial and macrophage differentiation potential

We next studied the differentiation capacity of aortic progenitors to determine whether they can contribute to the postnatal renewal of embryonically derived macrophages and endothelium. Progenitors, macrophages and endothelial cells were FACS-isolated from freshly digested 12 w C57BL/6J aortas and cultured in Matrigel^TM^ with endothelial growth medium. Whereas primary aortic macrophages displayed no cord forming activity, endothelial cells produced branching cord structures by day 7 (**Extended Data Fig. 8A**), with flow cytometry verifying their content to be CD45^−^CDH5^+^ endothelial cells (**Extended Data Fig. 8B**). Meanwhile, donor-matched aortic progenitors formed clusters by day 3, which sprouted and gave rise to complex cord networks by day 7, showing different morphology but similar overall cord length to those formed by primary endothelial cells (**Extended Data Fig. 8A**). Flow cytometry analysis of these networks showed that progenitors had produced a mixture of new CD45^+^CD11b^+^F4/80^+^ macrophages and CD45^−^CDH5^+^ endothelial cells, with some remaining progenitors (**Extended Data Fig. 8C**).

This assay was then repeated using progenitors isolated from adult aortic CFU-M. Culture-derived progenitors produced similar cord networks as freshly sorted progenitors (**Extended Data Fig. 8D**). In most donor mouse replicates, these contained a mixture of newly formed endothelial cells and macrophages as determined by flow cytometry (**Extended Data Fig. 8E, F**). In addition to losing surface expression of CD45 and gaining CDH5, the endothelial progeny of progenitors also acquired TIE2 (~80%) and VEGFR2 (~40%) expression (**Supplementary Table 1**). Meanwhile, the macrophages produced were mostly CX_3_CR1^+^ and CD206^+^ (~75%), with ~45% expressing LYVE-1 and only ~5% MHCII or CCR2 (**Supplementary Table 1**). We also labeled cytospin preparations of these cord networks with antibodies against EMCN and CD68 for confocal microscopy. This verified that progenitors had produced both EMCN^+^ endothelial cells and CD68^+^ macrophages, further supporting their bipotency (**Extended Data Fig. 8G**).

The bipotent capacity of aortic CFU-M progenitors was then studied at the clonal and single-cell levels. First, we harvested and disaggregated individual CFU-M and replated their cellular content in separate wells in Matrigel^TM^ for 7 d. Progenitors from single CFU-M formed sprouting networks, which contained both macrophages and endothelial cells (**Fig. 4A**), demonstrating that bipotency was contained at the clonal level in each of the five tested colonies. We then performed single-cell differentiation experiments. CFU-M were grown from adult ubiquitous GFP (UBI-GFP) (GFP^+^) and C57BL/6J (GFP^−^) aortas, individually isolated and disaggregated into single-cell suspensions. We seeded a single GFP^+^ progenitor with pooled GFP^−^ progenitors in the same well and co-cultured in Matrigel^TM^ for 7 d. Fluorescence microscopy showed formation of multicellular GFP^+^ sprouts from single-cell origins in multiple replicate experiments (**Fig. 4B**). Cytospin preparations of each well were labeled for GFP, EMCN and CD68. Five of eleven wells contained both GFP^+^EMCN^+^ endothelial cells and GFP^+^CD68^+^ macrophages (**Fig. 4B** and **Extended Data Fig. 9A,B**), while three had only GFP^+^EMCN^+^ cells and another three only GFP^+^CD68^+^ cells. This suggests that aortic CFU-M progenitors can differentiate into both endothelial and macrophage lineages at the single-cell level.

**Fig. 4.**
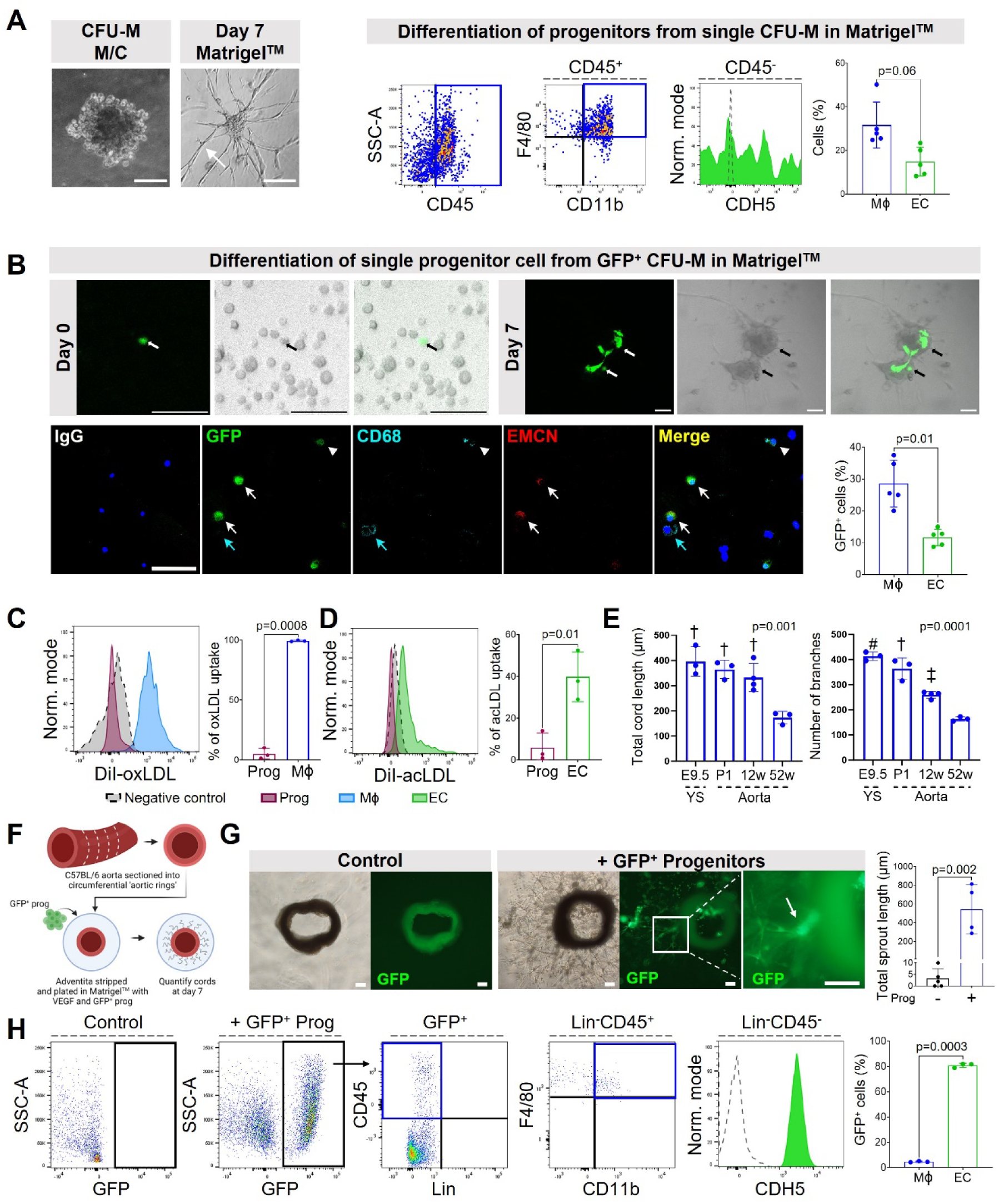
Aortic CFU-M progenitors have endothelial and macrophage potential. (A) Light microscopy images show a single CFU-M from C57BL/6J aorta and a branching network formed by its progenitors in Matrigel^TM^ after 7 d. Flow cytometry plots and graph show these networks contained new endothelial cells (EC) and macrophages (Mϕ) (n=5; Wilcoxon signed-rank test). (B) Top row: Confocal microscopy images (fluorescence, phase contrast and merged) show single GFP^+^ progenitor cell seeded with GFP^−^ progenitors at day 0 and resulting GFP^+^ sprout at day 7. Bottom row: Confocal microscopy images of immunolabeled cells show presence of GFP^+^CD68^+^ macrophage (arrowhead) and GFP^+^EMCN^+^ endothelial cells (white arrows). Cyan arrow indicates GFP^−^CD68^+^ macrophage. Merged image from corresponding IgG isotype negative control staining is also shown. *Please see Source Data file for larger version of these images.* Graph summarises frequency of macrophages and endothelial cells arising from a single GFP^+^ progenitor cell in replicate wells containing both cell types (n=5; paired t-test). (C, D) Uptake of (C) DiI-oxLDL or (D) DiI-acLDL by aortic progenitors and their macrophage or endothelial cell progeny produced in Matrigel^TM^ (n=3; paired t-tests). (E) Total cord length and number of branches produced in Matrigel^TM^ by E9.5 YS or aortic CFU-M progenitors from C57BL/6J mice of different ages (n=3-4; one-way ANOVA). Multiple test comparisons: †p<0.01, ‡p<0.001 and #p<0.0001 for comparison to 52 w. (F) Schematic of adventitial sprouting assay performed by culturing aortic progenitors from adult GFP mice with adventitia-less aortic rings from adult C57BL/6J mice. (G) Light and fluorescence microscopy images show adventitial sprouting without (control) and with (+) addition of GFP^+^ aortic progenitors, including higher magnification image of the inset box. Graph shows quantitative results for sprout length (n=4-5; unpaired t-test. (H) Flow cytometry plots and graph show the cells produced by culturing GFP^+^ aortic progenitors in aortic ring assay for 7 d (n=3; paired t-test). Data summarized as mean±SD. Scale bar, 100 µm. Also see **Extended Data Fig. 8** and **9** and **Supplementary Table 1**.

Macrophage and endothelial transformation of progenitors was also supported by new capacity of these progeny to take up oxidized and acetylated low-density lipoprotein (LDL) cholesterol, respectively, which was not a property of progenitors themselves (**Fig. 4C**,**D**). As was the case for 2° renewal, the cord-forming capacity of CFU-M progenitors was highest from E9.5 YS and P1 aortas and lower from 12 w and 52 w aortas (**Fig. 4E**).

We next examined whether aortic CFU-M progenitors mediate adventitial neovascularization, as occurs during *vasa vasorum* expansion. Angiogenic sprouting assays were performed with rings from ascending thoracic aorta of E9.5 TAM-induced adult *Cx3cr1*^CreER-YFP^ x *Rosa*^tdTom^ mice. These produced embryonically derived tdTom^+^ sprouts (**Extended Data Fig. 10A,B**), with a higher proportional content of tdTom^+^ cells compared to whole aorta, as measured by flow cytometry (**Extended Data Fig. 10C**). tdTom^+^ sprouts contained endothelial cells, progenitors and macrophages in decreasing order of abundance (**Extended Data Fig. 10D**).

We then used aortic rings from adult C57BL/6J mice, from which adventitia had been removed to eliminate sprouting (**Fig. 4F**). Seeding them with aortic CFU-M progenitors from adult UBI-GFP mice rescued adventitial angiogenesis with formation of GFP^+^ sprouts, that contained an abundance of endothelial cells, with a small percentage of macrophages that were predominantly LYVE-1^+^MHCII^−^ (**Fig. 4G,H** and **Supplementary Table 1**). Together, these data indicate that CFU-M progenitors participate in adventitial neovascularization.

To study the fate and vasculogenic capacity of CFU-M progenitors *in vivo*, we performed adoptive cell transfer experiments. Surgery was performed on 12 w C57BL/6J mice to induce hindlimb ischemia, before injecting the quadriceps and gastrocnemius with CFU-M-derived progenitors from adult aorta or E9.5 YS from UBI-GFP mice (~1.5×10^4^ cells) or cell-free Matrigel^TM^ (**Fig. 5A**). Laser Doppler imaging showed that aortic progenitors improved perfusion recovery over 14 d compared to control (**Fig. 5B**), accompanied by increased capillary and arteriolar density in injected muscle (**Extended Data Fig. 10E**). At day 14, GFP^+^ cells were detected in recipient muscle but not peripheral blood, with flow cytometry revealing that donor progenitors had produced new endothelial cells and macrophages (**Fig. 5C,D**). ~80% of the endothelial cells produced by progenitors *in vivo* were VEGFR2^+^, while most macrophages were again CX_3_CR1^+^LYVE-1^+^MHCII^−^, consistent with the characteristic surface marker profile described for embryonically derived tissue-resident macrophages ^14, 15^ (**Supplementary Table 1**). Confocal microscopy of immunolabeled sections also identified host-perfused, GFP^+^ endothelial-lined neovessels, with adjacent clusters of GFP^+^CD68^+^ macrophages (**Fig. 5E**). Similar results for perfusion recovery and endothelial and macrophage fate were obtained after injection of YS progenitors (**Extended Data Fig. 10F-H**).

**Fig. 5.**
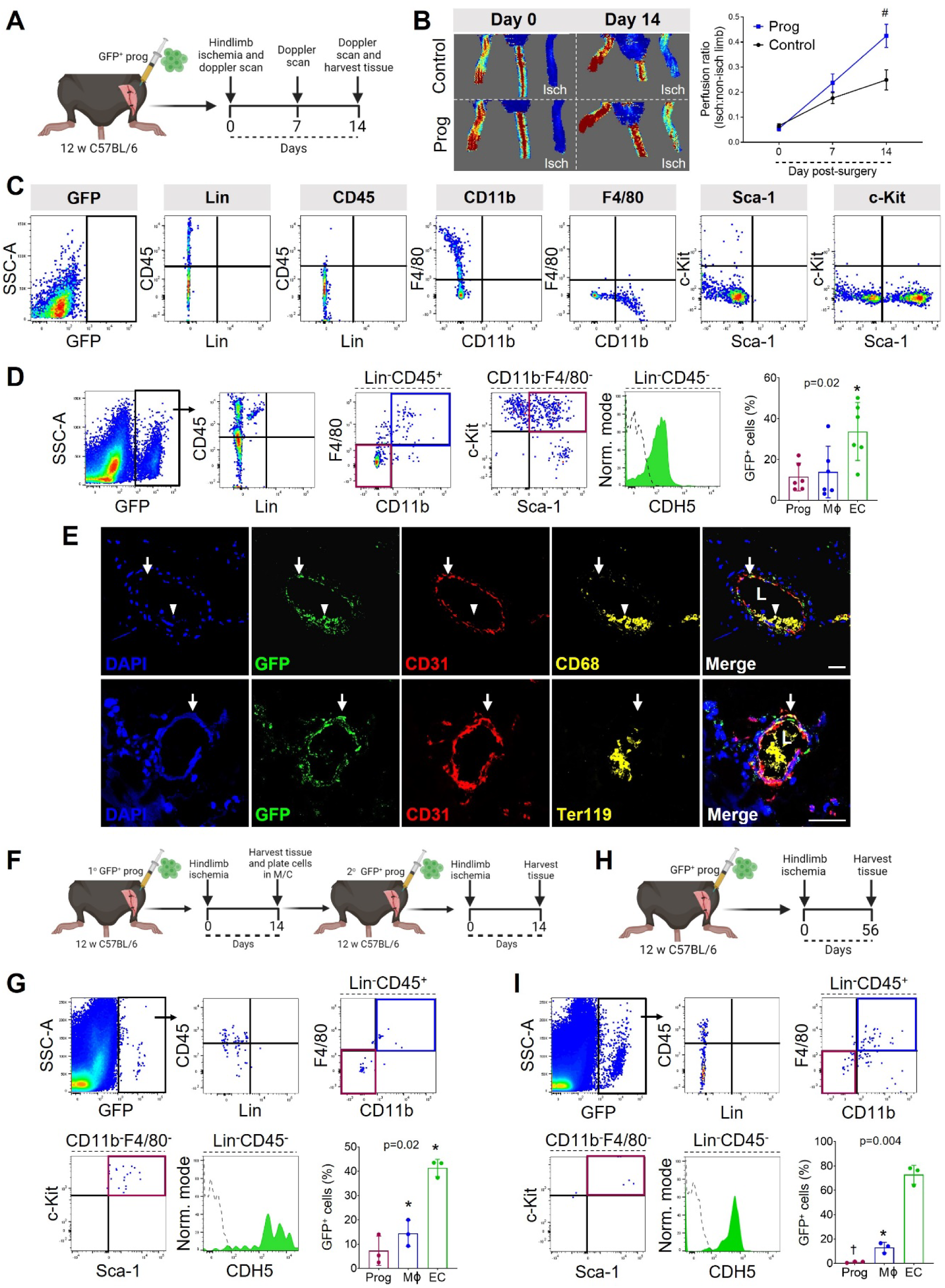
CFU-M progenitors have endothelial-macrophage plasticity and vasculogenic capacity *in vivo*. (A) Schematic of 1° transfer of GFP^+^ aortic progenitors in hindlimb muscle after hindlimb ischemia surgery with two-week follow-up. (B) Laser Doppler perfusion images of mice on day 0 and 14 after ischemia surgery and receiving cell-free control (above) or progenitors (below). Graph shows results over time (n=6/group). Mixed effects two-way ANOVA: p<0.0001 for time; p=0.03 for group; p=0.005 for time x group. Multiple comparison test: #p<0.0001 for progenitors vs control. (C) FMO control staining used to analyze the composition of cells produced by engrafted GFP^+^ progenitors after transfer into ischemic hindlimb muscle. (D) Flow cytometry plots and graph show the cells produced by donor cells in recipient muscle (n=6). Green histogram, sample; dotted histogram, FMO control. Repeated measures ANOVA. Multiple comparison test: *p<0.05 vs macrophages. (E) Confocal microscopy of immunolabeled recipient ischemic muscle shows neovessels lined by GFP^+^CD31^+^ endothelial cells (arrows) with cluster of GFP^+^CD68^+^ macrophages (arrowhead) (above) and perfused with host TER-119^+^ erythrocytes (below). L, lumen. (F) Schematic of 2° transfer of GFP^+^ progenitors into hindlimb muscle after hindlimb ischemia surgery. (G) Flow cytometry plots and graph show cells produced by donor cells after 2° transfer (n=3; repeated measures ANOVA). Multiple comparison test: *p<0.05 vs progenitors. (H) Schematic of 1° transfer of GFP^+^ aortic progenitors in hindlimb muscle after induction of hindlimb ischemia with 8 w follow-up. (I) Flow cytometry plots and graph show cells produced by donor cells 8 w after 1° transfer (n=3; repeated measures ANOVA). Multiple comparison test: *p<0.05 and †p<0.01 vs endothelial cells. Data summarized as mean±SD. Scale bar, 20 µm. Also see **Extended Data Fig. 10**.

Collectively, the above results identify aortic CFU-M progenitors as vasculogenic EndoMac progenitors. We next examined their renewal capacity and the durability of their progeny *in vivo*. Progenitors from adult UBI-GFP aortas were transplanted into ischemic C57BL/6J hindlimbs as above. 14 d later, we digested the recipient quadriceps and gastrocnemius into single-cell suspensions that were plated in methylcellulose for another 14 d. These generated GFP^+^ CFU-M in methylcellulose, from which GFP^+^ progenitors were again isolated and used in 2° hindlimb transfer studies (**Fig. 5F**). After another 14 d, engrafted GFP^+^ cells had again transformed into endothelial cells or macrophages, with a small percentage of residual progenitors (**Fig. 5G**). Similar results were obtained in 1° transfer studies that were followed for eight weeks instead of two (**Fig. 5H,I**). These results show that EndoMac progenitors from adult aorta produce durable endothelial and macrophage progeny *in vivo*. Although they do renew, their capacity to do so is finite meaning that their own numbers diminish over time.

### EndoMac progenitors exhibit a myelopoietic and vasculogenic transcriptional profile

To further examine the cellular composition of progenitors contained within adult aortic CFU-M, we performed single cell RNA sequencing (scRNA-seq) of culture-derived progenitors pooled from two 12 w C57BL/6J mice. After quality control and filtering, transcriptional profiles of 7,966 cells were analyzed and expression of a total of ~26,000 genes was detected (**Supplementary Table 2**). Cluster analysis revealed that CFU-M progenitors were composed of nine clusters of cells, including six closely related clusters (progenitors 1-6), two highly proliferative clusters marked by *Mki67* expression (Prolif progenitors 1 and 2), and a small distinct cluster (progenitors 7) (**Fig. 6A-D** and **Supplementary Table 3**). Consistent with the immunophenotype of aortic CFU-M progenitors, all clusters expressed *Ptprc* (CD45), *Kit*, *Ly6a* (Sca-1) and *Pecam1* (CD31), with almost no expression of mature endothelial (*Cdh5, Kdr/*Vegfr2, *Tek/*Tie2) or macrophage (*Itgam/*Cd11b, *Adgre1/*F4/80, *Fcgr1/*Cd64, *Lyve1*) genes (**Fig. 6E,F**). Near absence of *Flt3* expression reiterated that these cells originate independently of Flt3-mediated hematopoiesis (**Fig. 6G**), while expression of *T* (Brachyury) in most cells indicated their origin from embryonic mesenchyme (**Fig. 6H**). In keeping with our earlier results, the progenitors did not express pluripotency genes (*Nanog, Sox2, Pou5f1/Oct4*). However, all the clusters expressed the stem cell self-renewal gene *Klf2*, six of the clusters also expressed low levels of *Myc* and in addition, the smallest cluster (progenitors 7) expressed *Klf4* (**Extended Data Fig. 11A**). Genes and transcription factors that mark YS EMPs were expressed in the majority of cells (*Cd93*, *Gata1*, *Gata2*) or at least in some cells in all clusters (*Csf1r*, *Bpnt1*, *Id1*), consistent with their embryonic YS origin (**Fig. 6I** and **Extended Data Fig. 11B**) ^43^. At the same time, all clusters displayed variable levels of expression of *Cx3cr1*, *Zeb2*, *Maf*, *Mrc1* and *Tnfrsf1b* genes, which are upregulated in YS-derived pre-macrophages ^43^ (**Fig. 6I** and **Extended Data Fig. 11C**). Additionally, albeit at varying frequencies, all clusters expressed many of the genes involved in myelopoiesis (*Spi1, Cd34, Myb, Mafb, Cebpb, Nr4a1*) ^44–48^ and vasculogenesis or angiogenesis (*Gsn, S100a6, Dcn, Runx1, Vegfa, Flt1, Angpt1, Epor*) ^49–51^, while cluster 7 was notable for expressing higher levels of *Flt1*, *Il33, Pdgfra, Postn, Cd248* and *Sox9*, which are upregulated in endothelial progenitor cells ^51–53^ (**Fig. 6I** and **Extended Data Fig. 11D,E**). This, in line with our *in vitro* and *in vivo* findings, suggests that CFU-M progenitors are transcriptionally primed for differentiation into myelopoietic and vasculogenic lineages.

**Fig. 6.**
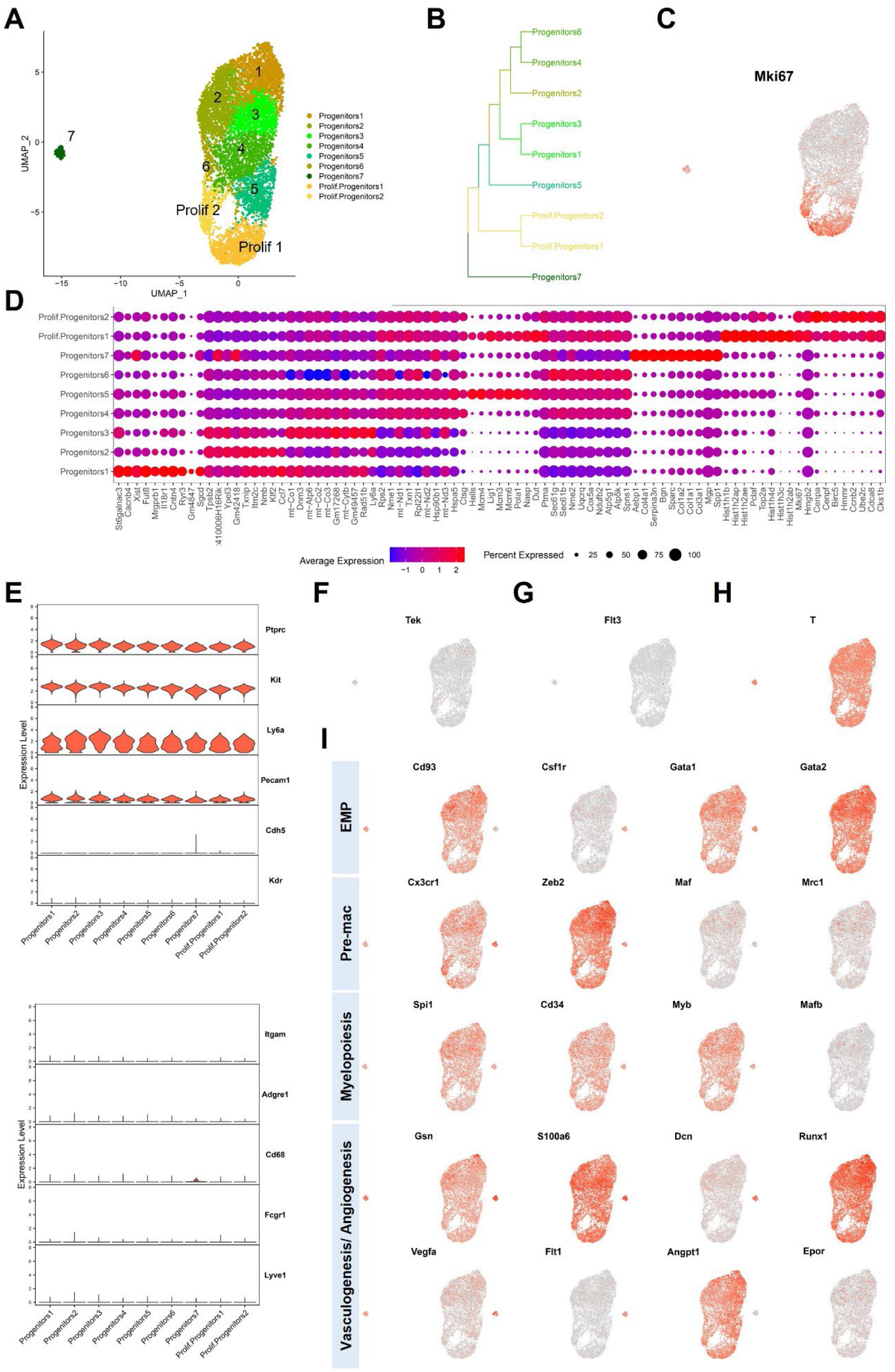
Aortic CFU-M progenitors exhibit a myelopoietic and vasculogenic transcriptional profile. Aortic CFU-M progenitors cultured from two adult C57BL/6J mice were pooled and viable cells analyzed by scRNA-seq. (A) UMAP plot of 7,966 cells that passed quality controls colored by cluster assignment. Each dot represents a cell. Cell clusters are colored as indicated. (B) Phylogenetic tree shows the hierarchical relationships of assigned clusters. (C) Expression levels of *Mki67* overlaid on the UMAP plot showing cell specificity of expression. Expression levels are shown as log normalised counts. The grey to red gradient represents low to high values. Predominant expression can be seen in the proliferative clusters. (D) Dot plot shows the top 10 differentially expressed genes in each cluster and relative expression levels of those genes in all clusters. Purple to red gradient represents average expression levels from low to high. Dot size denotes percentage of cells expressing a gene, as indicated. (E) Violin plots illustrate cluster-wise expression of the genes corresponding to the key surface markers expressed by progenitors, along with selected mature endothelial (top) and macrophage marker (bottom) genes. F) Expression levels of the endothelial gene *Tek* overlaid on the UMAP plot showing minimal expression. (G-I) Expression levels of the indicated marker genes for G) definitive hematopoiesis, H) hemangioblasts from embryonic mesenchyme, I) YS EMPs (EMP), pre-macrophages (Pre-mac), myelopoiesis and vasculogenesis/angiogenesis, overlaid on the UMAP plot showing cell specificity of expression. Also see **Extended Data Fig. 11**.

Trajectory analysis using Monocle3 revealed a progressive relationship between clusters of cells in pseudotime starting from cluster 1 and ending in proliferative clusters, indicating that they represent different cell states (**Extended Data Fig. 11F**). Pathway analysis of gene expression signatures of cellular clusters (**Supplementary Table 4**) showed an overlap of overrepresented pathways among clusters, also suggesting a continuum of cell states (**Extended Data Fig. 11G** and **Supplementary Tables 5-13**). Transcription factor binding site analysis of gene expression signatures revealed significant overrepresentation of binding sites for transcription factors involved in the regulation of cell cycle (members of E2F and DP families and Rb), embryonic development (Foxd3 and Sox10), cell fate determination (Sox10), genes in response to injury and inflammation (Nrf2), heart development (Zfp161) and circulatory system development (Hes1), in the differentially expressed genes in one or more clusters (**Supplementary Table 14**). This is consistent with the tissue source, developmental origin, and properties of these progenitors.

Together, these data demonstrate that aortic CFU-M progenitors are a relatively homogeneous population containing highly proliferative as well as less proliferative cells with transcriptional priming for myelopoiesis and vasculogenesis, and some degree of heterogeneity that is likely attributable to asynchronous cell states.

### Regulatory effects of Angiotensin II on aortic EndoMac progenitors

The renin-angiotensin system, and specifically AngII, play key regulatory roles in myelopoiesis ^54^ and vascular inflammation ^55^. When pluripotent stem cells undergo hemato-endothelial differentiation *in vitro*, they transition through bipotent hemangioblasts that form colonies and express ACE (CD143) and the receptors for AngII, AGTR1 and AGTR2 ^31^. We found that ACE was expressed on EndoMac progenitors from adult aortic CFU-M *in vitro*, adult aorta *in vivo* and E9.5 YS CFU-M (**Fig. 7A-C**). Aortic progenitors also expressed AGTR2 at the cell surface (**Fig. 7A**) and at mRNA level (**Fig. 7D**), with much lower expression of AGTR1. AngII induced concentration-dependent increases in 1° CFU-M yield from 12 w C57BL/6J aortas (**Fig. 7E**). Furthermore, when incubated with 100 nM AngII, aortic CFU-M could be renewed in bulk culture for four passages in much larger numbers compared to control (**Fig. 7F**). Conversely, treatment with inhibitors of ACE (enalapril), AGTR1 (losartan) and AGTR2 (PD123,319) showed that inhibition of AGTR1 and AGTR2 reduced 1° CFU-M yield (**Fig. 7G**).

**Fig. 7.**
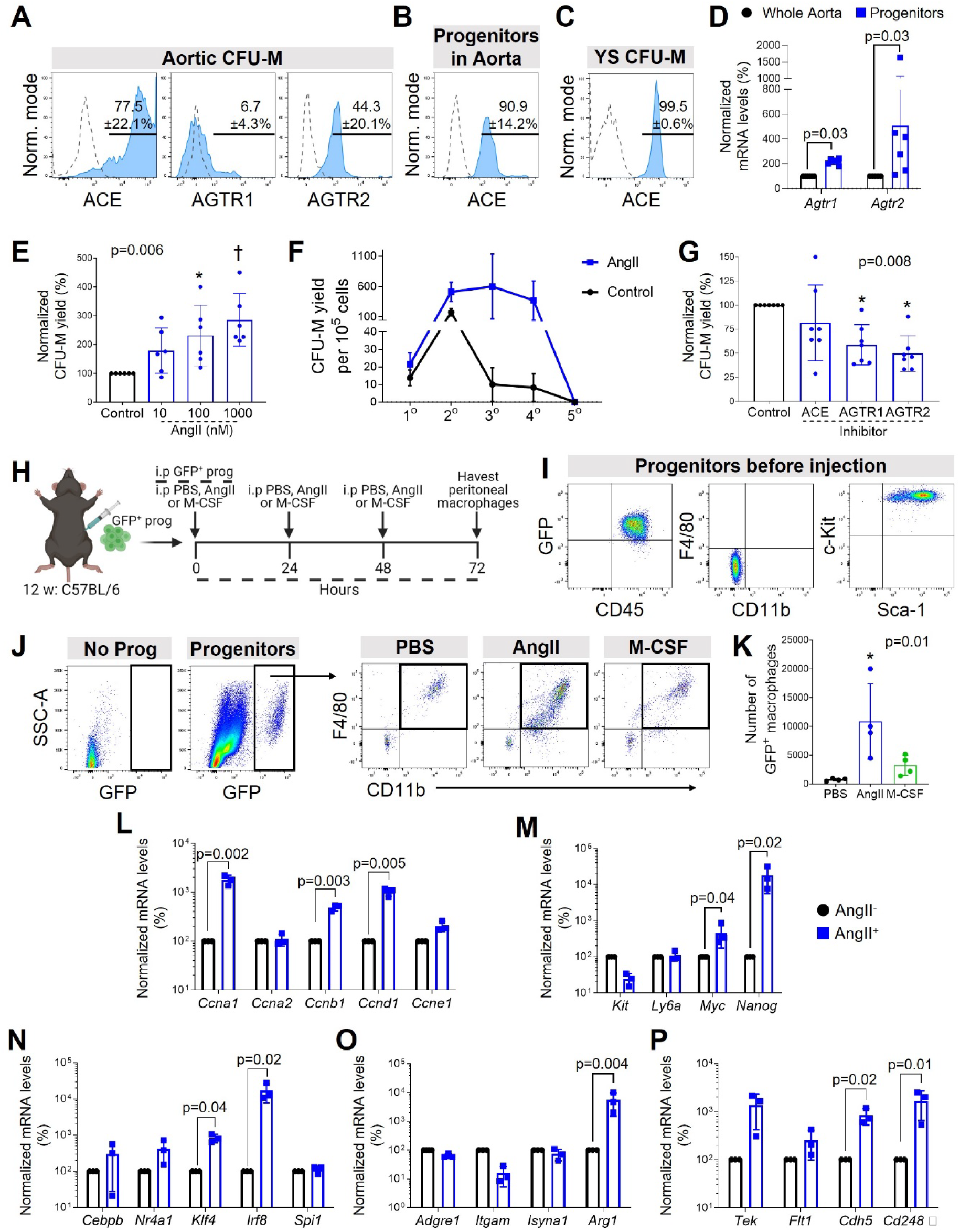
Regulatory effects of Angiotensin II on aortic EndoMac progenitors. (A) Surface expression of ACE, AGTR1 and AGTR2 on progenitors from aortic CFU-M (n=5-6 12 w C57BL/6J mice). (B) ACE expression on progenitors in 12 w C57BL/6J aortic digests (n=4). (C) ACE expression on progenitors from E9.5 C57BL/6J YS CFU-M (n=3). (D) Normalized mRNA expression of *Agtr1* and *Agtr2* in progenitors from aortic CFU-M relative to donor-matched aortic digests (n=6 12w C57BL/6J mice). Wilcoxon signed-rank tests. (E) Aortic CFU-M yield with different concentrations of AngII, normalized to no AngII control (n=6 12 w C57BL/6J mice). Friedman test. Multiple comparison tests: *p<0.05, †p<0.01 vs control. (F) Aortic CFU-M yield across serial passages in the presence or absence of 100 nM AngII (n=3-4 12 w C57BL/6J mice). (G) Aortic CFU-M yield in the presence of inhibitors of ACE (Enalapril), AGTR1 (Losartan) and AGTR2 (PD123319) normalized to no inhibitor control (n=7). Friedman test. Multiple comparison tests: *p<0.05 vs control. (H) Schematic of peritoneal transfer assay. GFP^+^ aortic progenitors were intraperitoneally injected into 12 w C57BL/6J mice with daily injections of PBS, AngII or M-CSF for 72 h. (I) Immunophenotype of GFP^+^ aortic progenitors before intraperitoneal injection. (J) Flow cytometry shows *de novo* formation of macrophages from GFP^+^ aortic progenitors in peritoneal cavity after 72 h. No progenitor cells (Prog) negative control for GFP also shown. (K) Number of macrophages produced by progenitors under different conditions (n=4; one-way ANOVA). Multiple comparisons test: *p<0.05 vs PBS. (L-P) mRNA expression of selected genes in aortic progenitors after treatment with AngII. Expression normalized to *β-actin* and then no AngII control. Genes relate to (L) cell cycle; (M) progenitor/stem cell biology and self-renewal; (N) myelopoiesis; (O) macrophages; (P) endothelial biology and angiogenesis (n=3 12 w C57BL/6J mice; paired t-tests). Data summarized as mean±SD.

Given that AngII rapidly stimulates the accumulation of macrophages in aortic adventitia ^56^, we next studied how it affects the ability of aortic progenitors to produce macrophages *in vivo*. We made use of the peritoneal cavity as a niche for macrophage expansion by injecting 10^4^ progenitors from 12 w UBI-GFP aortas into the peritoneum of C57BL/6J mice, and followed this with daily *i.p.* injections of AngII, M-CSF or PBS (**Fig. 7H**). Flow cytometry was used to check the purity of undifferentiated progenitors before injection (**Fig. 7I**) and their *de novo* differentiation into CD11b^+^F4/80^+^ macrophages on retrieval from the peritoneal cavity 72 h later (**Fig. 7J**). AngII promoted expansion of progenitor-derived macrophages by 14.5-fold and 3.4-fold compared to PBS and M-CSF, respectively (**Fig. 7J,K**).

In keeping with these different effects of AngII on EndoMac progenitors, qRT-PCR also showed that it upregulated mRNA levels of genes involved in cell division (*Ccna1*, *Ccnb1*, *Ccnd2*), self-renewal (*Myc*, *Nanog, Klf4*) ^47, 57^, myelopoiesis (*Klf4*, *Irf8*) ^58^, M2-like macrophage polarization (*Arg1*) and endothelial specification and angiogenesis (*Cdh5*, *Cd248*) (**Fig. 7L-P**).

### EndoMac progenitors expand early in response to AngII-induced aortic inflammation

AngII-induced vascular inflammation is characterized by expansion of adventitial macrophages ^56^, adventitial fibrosis ^59^ and in some mouse strains, development of aortic aneurysms ^60^. Another study reported that after 10 d of exposure to AngII, adventitial macrophages expanded due to both recruitment of BM-derived cells and local proliferation of embryonically derived macrophages ^36^. As AngII stimulated a marked increase in progenitor-derived macrophages within 72 h in our peritoneal transfer assay, we focused on its early effects on aortic progenitors and macrophages after systemic infusion by osmotic pump. In *Flt3*^Cre^ x *Rosa*^mT/mG^ mice, 48 h of AngII infusion caused a significant 5.8-fold increase in Flt3-Cre^−^ macrophages compared to PBS control (**Fig. 8A,B**), with more Flt3-Cre^+^ and Flt3-Cre^−^ macrophages in S-phase of cell cycle (both 2.9-fold vs PBS) (**Fig. 8C**). As a potential contributor to the expansion of non-BM derived macrophages, AngII also induced expansion and proliferation of Flt3-Cre^−^ progenitors, with fold comparisons compared to PBS: 3.0x for aortic CFU-M yield (**Fig. 8D**); 8.5x for progenitor number assessed by flow cytometry (**Fig. 8E**); and 10.5x for progenitors in S-phase (**Fig. 8F**).

**Fig. 8.**
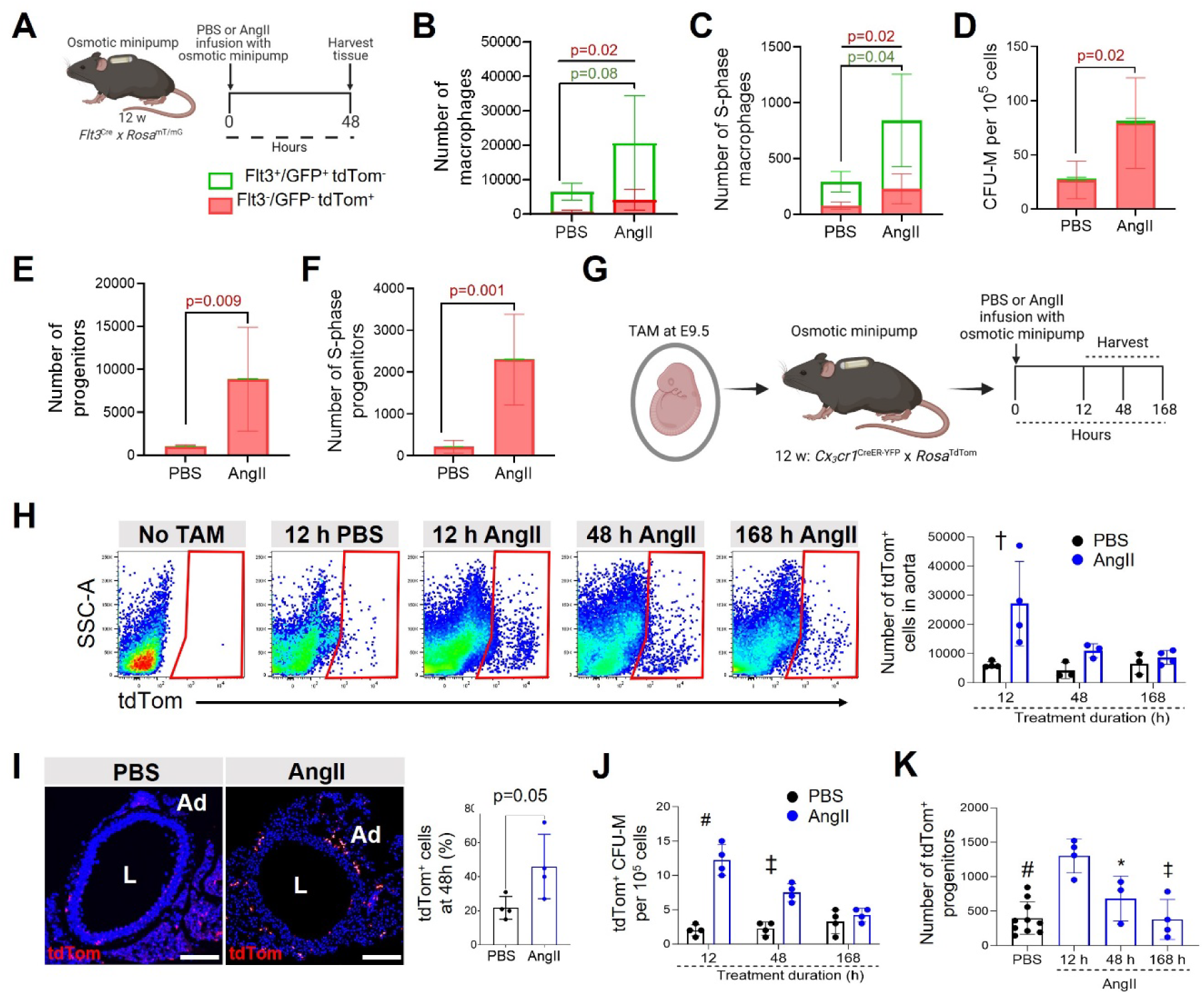
Angiotensin II-induced vascular inflammation involves early expansion of EndoMac progenitors *in vivo*. (A) Schematic of PBS or AngII infusion in adult *Flt3*^Cre^ x *Rosa*^mT/mG^ mice for 48 h. (B-F) Comparisons of aortas from PBS and AngII groups for the numbers of Flt3-cre^+^ (green) and Flt3-cre^−^ (red): (B) macrophages, (C) S-phase macrophages, (D) CFU-M yield, (E) progenitors and (F) S-phase progenitors (n=5-6/group). Flow cytometry used for (B), (C), (E) and (F). Unpaired t-tests. P-values in red, Flt3-cre^−^ comparisons; in green, Flt3-cre^+^ comparisons. (G) Schematic of PBS or AngII infusion in E9.5 TAM-induced 12 w *Cx3cr1*^CreER-YFP^ x *Rosa*^tdTom^ mice with aortas harvested at different times. (H) Flow cytometry of tdTom^+^ labeling in aortic digests at different times. No TAM control also shown. Graph summarizes number of tdTom^+^ cells in aortas (n=3-4). Two-way ANOVA: Interaction p=0.04, Time p=0.005, Group p=0.03. Mann-Whitney or unpaired t-tests also used to compare groups for each time point: †p<0.01. (I) Confocal microscopy of immunolabeled sections of descending aorta after 48 h of PBS or AngII. Graph shows % of tdTom^+^ cells in adventitia (Ad) (n=4, three sections each). Unpaired t-test. L, lumen. Nuclei stained with DAPI (blue). (J) tdTom^+^ aortic CFU-M yield (n=4). Two-way ANOVA: Interaction p<0.0001, Time p=0.0005, Group p<0.0001. Unpaired t-tests also used to compare groups for each time point: #p<0.0001, ‡p<0.001. (K) Numbers of tdTom^+^ progenitors in aorta after different durations of AngII or PBS (n=3-10). One-way ANOVA p=0.0001. Multiple comparisons test: *p<0.05, ‡p<0.001, #p<0.0001 vs 12 h. Data summarized as mean±SD. Scale bar, 50 μm.

We next used adult *Cx3cr1*^CreER-YFP^ x *Rosa*^tdTom^ mice that had been administered TAM at E9.5, for AngII or PBS infusion by osmotic pump for 12 h, 48 h or 168 h (**Fig. 8G**). AngII increased the numbers of tdTom^+^ cells in aorta at 12 h and 48 h compared to PBS (fold difference by flow cytometry: 4.4x at 12 h, 2.6x at 48 h, 1.3x at 168 h) (**Fig. 8H**). Confocal microscopy of immunolabeled aortic sections showed a similar expansion of tdTom^+^ cells in the adventitia of descending aorta at 48 h (**Fig. 8I**). As measured by aortic CFU-M yield, the AngII-induced expansion of embryonically derived EndoMac progenitors was highest at 12 h (6.1-fold vs PBS), still evident at 48 h (3.3-fold) but no longer at 168 h (1.3-fold) (**Fig. 8J**). This was corroborated by flow cytometry, showing that EndoMac progenitors undergo very early expansion in response to AngII (**Fig. 8K**).

## DISCUSSION

The intimate association between the hematopoietic and endothelial lineages begins early in development. Converging evidence indicates that YS EMPs give rise to both macrophages and endothelial cells in adult mouse tissues ^16, 28^. In this study we identify undifferentiated EndoMac progenitors in murine aorta that also originate embryonically. These progenitors are more abundant at birth, at which time they have enhanced clonal capacity and are maintained independently of Flt3^+^ BM hematopoiesis. As supported by *in vitro* and *in vivo* differentiation studies, this population is also bipotent for macrophage and endothelial lineages and possesses vasculogenic capacity. We also demonstrate a regulatory role for AngII, which stimulates its clonogenic, proliferative and differentiation properties.

Most mature cell types rely on the self-renewal of stem cells and proliferation and differentiation of transient amplifying progenitors for their homeostatic turnover and recovery after tissue insult. Although there has been conjecture about how embryonically derived macrophages are maintained after birth, prevailing opinion favors their ability to self-renew without losing functional or differentiated status ^27^. This was initially based on experiments showing self-renewal of macrophages in which MafB and c-Maf were genetically knocked out ^46^. Subsequent studies have provided contrasting data either supporting the long-lived nature of tissue-resident macrophages, especially microglia ^61^, their stochastic turnover ^13, 62^, repopulation by clonal expansion ^63, 64^ or differentiation from local progenitors ^65^. The mechanistic basis for macrophage self-renewal has been linked to downregulation of MafB/cMaf, which occurs constitutively in some macrophage populations (e.g. alveolar macrophages) or can be induced (e.g. by M-CSF). This in turn activates a self-renewal gene network, centered around *Myc*, *Klf2* and *Klf4* ^47^.

As shown here, the *in vitro* growth of CFU-M from adult aorta is not due to monocytes or macrophages, but rather embryonically derived progenitors. The finding that CFU-M undergo 2° renewal from single-cell origins paved the way for us to study their composition and identify a Lin^−^CD45^+/Lo^CD11b^−^F4/80^−^Sca-1^+^c-Kit^+^ fingerprint for progenitors both in culture-derived CFU-M and aorta *in vivo*. This surface phenotype distinguishes EndoMac progenitors from myeloid-committed progenitors in BM, such as macrophage/dendritic cell progenitors and common monocyte progenitors, which lack Sca-1 but express Flt3/CD135 and Ly6C, respectively ^66, 67^. In prior studies we found that BM cell transplantation made minimal contribution to the recovery of CFU-M progenitors in aortic adventitia after depletion by whole body irradiation, and the recovery of progenitors was slower than blood cells, which indicated their non-BM hematopoietic origin ^34, 35^. This is supported by the present study that shows that EndoMac progenitors are largely independent of Flt3-mediated hematopoiesis. However, given that Flt3-Cre labeling efficiency was incomplete in the *Flt3*^Cre^ x *Rosa*^mT/mG^ model used here, we cannot rule out that a minority of progenitors could still have Flt3^+^ origins, or alternatively could directly arise from Flt3^−^ BM LT-HSCs by circumventing differentiation via ST-HSCs and MPPs, which requires further investigation. As a high percentage of progenitors are actively dividing even in steady state, it is not surprising that their ability to renew is finite and diminishes with aging, along with their clonogenic and angiogenic capacity. Combined with a lack of circulatory renewal from Flt3^+^ BM hematopoiesis, this helps explain why aortic progenitor numbers decrease postnatally. However, we also demonstrate that their proliferation and renewal can be amplified by stimulatory cues, such as elicited by AngII.

As in previous studies ^68^, we found negligible contribution from YS EMPs to adult BM hematopoietic cells. As revealed by embryonic profiling across different gestational ages and timed induction of fate-mapping, EndoMac progenitors are present in YS by E8.5 and migrate intra-embryonically to AGM by E10.5. This aligns with prior findings for EMPs and CX_3_CR1^+^ pre-macrophages ^24^. Throughout our differentiation assays, EndoMac progenitors produced macrophages without evidence of passing through a monocyte intermediate stage. This separates them from the EMPs that colonize fetal liver where they produce monocytes and other lineage-committed progenitors ^25^. This was reflected here by mixed growth of different CFU types from E12.5-15.5 liver, as distinct from selective CFU-M growth from AGM at the same ages. By scRNA-seq we found that culture-derived progenitors expressed genes transcribed in YS EMPs and pre-macrophages, which places them in an intermediate stage and may explain their ability to directly produce macrophages.

The classical origin of embryonic vascular endothelial cells is the differentiation of mesoderm-derived angioblasts around E7.0 YS in mice ^69^. Subsequently, endothelial cells undergo local proliferation during tissue angiogenesis, with evidence for clonal expansion in adult tissues under ischemic insult ^70^. Other identified sources of postnatal endothelial renewal include circulating endothelial progenitor cells ^71^ and tissue-resident endovascular progenitors ^52^. Plein *et al*. previously reported that endothelial cells in YS and some embryonic and adult tissues also derive from YS EMPs ^28^, although this was not reproduced by another study ^29^. In tracking the fate of E8.5 and 9.5 CX_3_CR1^+^ and E8.5 CSF1R^+^ progenitors, we examined both their macrophage and endothelial progeny. In support of Plein’s study, we identified embryonically derived endothelial cells in postnatal aorta.

Although it has been widely accepted that embryonically derived macrophages and endothelium are maintained after birth by proliferative self-renewal, another explanation could be the postnatal differentiation of EndoMac progenitors into these two distinct lineages. We observed this to occur across different *in vitro* and *in vivo* experiments with overrepresentation of implicated signaling pathways and expression of myelopoietic and vasculogenic genes *in vitro*. While our results suggest that these progenitors have a differentiation bias toward the endothelial lineage in the settings of adventitial vasculogenesis and post-ischemia repair, they can also rapidly generate macrophages, as seen here after peritoneal injection. Moreover, they appear predisposed to forming macrophages that are LYVE-1^+^MHCII^−/Lo^CCR2^−^, in keeping with the identity of embryonically derived macrophages that reside in the aortic adventitia and other tissues ^15, 36, 39^. Importantly, we established that CFU-M derived progenitors can produce both endothelial cells and macrophages in clonal and single-cell assays *in vitro*. Along with their relative transcriptional homogeneity by scRNA-seq, this suggests that their bipotency is contained at the single-cell level rather than being the byproduct of a mixed population of unipotent endothelial and macrophage progenitors. However, this requires definitive confirmation in future studies by using a genetic cell tagging approach.

The century-old notion of the hemangioblast, a bipotent progenitor for endothelial and hematopoietic cells in development ^30^, has been supported by *in vitro* evidence of a common pathway for hemato-endothelial differentiation from pluripotent stem cells ^31, 32, 72^. Here, we observed by scRNA-seq that EndoMac progenitors express the canonical hemangioblast gene, *T* (Brachyury), while they also exhibited surface expression of ACE and AGTR2, which too are linked to hemato-endothelial bipotency ^31^. Focusing our attention on AngII as a regulator of their properties, we found that AngII stimulated the proliferative, clonogenic and macrophage-forming capacity of aortic EndoMac progenitors, and upregulated various genes, such as *Nanog*, *Myc*, *Irf8* and *Arg1*. Together with our results from short-term AngII infusion in *Flt3*^Cre^ and *Cx3cr1*^CreER^ mice, this indicates that these progenitors are primed to provide an immediate proliferative response to AngII. This may help feed the local expansion of adventitial macrophages and endothelial cells, which is complemented by recruitment and proliferation of BM-derived macrophages, which also contribute to AngII-induced adventitial inflammation ^36^.

In conclusion, our discovery of aortic EndoMac progenitors adds to the recognized fate of early embryonic progenitors in postnatal tissues. These bipotent progenitors can rapidly proliferate and differentiate to drive adventitial neovascularization. Their existence also provides a new model to help explain the local maintenance and expansion of embryonically derived tissue-resident macrophages and endothelial cells after birth.

## METHODS

### Resource availability

Details of general reagents are provided in **Supplementary Table 15**. Further information and requests for resources and reagents should be directed to and will be fulfilled by the lead contact, Peter Psaltis.

### Experimental model details

#### Mouse models

*Cx3cr1*^GFP/+^ (B6.129P-Cx3cr1^tm1Litt^/J), *Ccr2*^−/−^ (B6.129S4-Ccr2^tm1Ifc^/J), *Rosa*^mTmG^ (Gt(ROSA)26Sor^tm4(ACTB-tdTomato,-EGFP)Luo^/J), *Csf1r*^Mer-iCre-Mer^ (FVB-Tg(Csf1r-cre/Esr1*)1Jwp/J), *Cx3cr1*^YFP-creER^ (B6.129P2(Cg)-Cx3cr1^tm2.1(cre/ERT2)Litt^/WganJ) and UBI-GFP (C57BL/6-Tg(UBC-GFP)30Scha/J) mice were purchased from The Jackson Laboratory. C57BL/6J mice were from the South Australian Health and Medical Research Institute (SAHMRI). Male *Flt3*^Cre^ breeding mice ^73^ were initially provided by Professors Thomas Boehm (Max-Planck-Institute of Immunobiology and Epigenetics, Freiburg, Germany) and Toshiaki Ohteki (Tokyo Medical and Dental University, Tokyo, Japan). A breeding colony of *Rosa*^tdTom^ (B6.Cg-*Gt(ROSA)26Sor*^tm14(CAG-tdTomato)Hze^/J) mice was provided by Dr Daniel Worthley (Precision Medicine Theme, SAHMRI).

*Cx3cr1*^GFP/+^ mice were inter-crossed to obtain *Cx3cr1*^GFP/GFP^ mice. *Flt3*^Cre^ and *Csf1r*^Mer-iCre-Mer^ mice were crossed with *Rosa*^mTmG^ mice to obtain *Flt3*^Cre^ x *Rosa*^mTmG^ and *Csf1r*^Mer-iCre-Mer^ x *Rosa*^mTmG^ mice, respectively. *Cx3cr1*^YFP-creER^ mice were crossed with *Rosa*^tdTom^ mice to obtain *Cx3cr1*^CreER-YFP^ x *Rosa*^tdTom^ mice. *Csf1r*^Mer-iCre-Mer^ mice were maintained on a FVB and C57BL/6J mixed background. All mice were housed in a temperature and humidity-controlled environment under a 12-hour light/dark cycle with *ad libitum* access to water and standard chow diet. Male and female mice were used in all experiments and experimental arms were gender- and age-matched, except for *Flt3*^Cre^ x *Rosa*^mTmG^ fate-mapping studies where only males were used as the Flt3-cre modification is located on the Y chromosome. All animals were treated in accordance with the Australian Code of Practice for the Care and Use of Animals for Scientific Purposes and were approved by the SAHMRI Animal Ethics Committee (ID SAM117, SAM155, SAM308 and SAM432.19).

To achieve Cre-Lox recombination in *Cx3cr1*^CreER-YFP^ x *Rosa*^tdTom^ and *Csf1r*^Mer-iCre-Mer^ x *Rosa*^mTmG^ mice, 75 µg/g of 4-hydroxytamoxifen (TAM, Sigma-Aldrich) was intraperitoneally (*i.p.*) injected into pregnant dams at either E8.5 or E9.5, as specified. To analyze the cell cycle state of progenitors in aorta, 1 mg of bromodeoxyuridine (BrdU) was injected *i.p.* before euthanasia and tissues harvested at specified times.

#### Preparation of single-cell suspensions

Experiments were performed with freshly isolated, single cell disaggregates from tissues, as specified. Aortas were dissected out intact, along the entire length from aortic valve to iliac bifurcation and flushed extensively with PBS, before microscopic dissection of surrounding perivascular fat and, where indicated, careful separation of the adventitia. The aorta was digested with Liberase^TM^ TM (50 μg/mL) (Roche Applied Science) for 1.5 h at 37°C. Tissue digests were then passed through a 40-µm nylon mesh (Greiner Bio-One) and neutralized with Iscove’s Modified Dulbecco’s Medium (IMDM, Sigma-Aldrich) supplemented with 10% fetal bovine serum (FBS, Cell Sera).

Where applicable, single cell disaggregates were also prepared from the liver, quadriceps and gastrocnemius muscle, brain, blood, BM and spleen. Liver and muscle tissues were digested with Liberase as per the aortas to give single-cell suspensions. The brain was carefully removed from cephalic mesenchyme and meninges, dissected out, minced, and incubated in Liberase^TM^ TM (50 μg/mL) for 2 h and passed through a 40-µm nylon mesh ^74^. Digests of brain were resuspended in 37% isotonic Percoll, underlaid with 70% isotonic Percoll (GE Healthcare), centrifuged (600 *g*, 25 min) and cells in the interface were collected. Blood was collected by cardiac puncture in EDTA coated blood collection tubes. Erythrocytes were lysed by mixing with ammonium chloride (1:10 v/v; StemCell Technologies) at 4°C for 10 min, then non-erythroid blood cells were washed and centrifuged (300 *g*, 5 min). BM cell suspensions were prepared by flushing femurs and tibias with PBS. Spleens were homogenized through a 40-μm cell strainer. BM and spleen cell suspensions were incubated in ammonium chloride at 4°C for 8 min, following which cells were washed and centrifuged (400 *g*, 5 min).

Gestational embryonic age was defined based of the date of vaginal plug formation, which was set at E0.5. To obtain single cell disaggregates from embryos, pregnant females were euthanized by CO_2_ exposure or cervical dislocation. Each embryo was carefully dissected from the uterus under a dissection light microscope (Carl Zeiss); tissues harvested were YS (E7.5-E15.5), whole embryo (E7.5-E9.5), AGM (E10.5-E15.5) and liver (E11.5-E15.5) ^75^. Tissues were washed with PBS and placed in Hanks balanced-salt solution (HBSS, Sigma) supplemented with 2% FBS and digested with 1 mg/mL of collagenase II for 15 min at 37°C. Tissue suspensions were filtered using Polystyrene Round-Bottom Tubes with Cell-strainer cap strainers (In Vitro Technologies) and centrifuged (400 *g*, 5 min).

All single-cell suspensions were resuspended in IMDM supplemented with 10% FBS and 1% antimycotic/antibiotic solution, before performing total cell counts for colony-forming unit (CFU) assays or flow cytometric characterization.

For some experiments, YS tissue was digested with collagenase II (273 U/ml) for 10-15 min at 37°C and digestion stopped with 100% FBS. Digested single cell disaggregates were filtered through a 40-µm mini cell strainer (Cell Systems Biology) and washed with PBS supplemented with 2% FBS. Cells were collected by centrifugation (300 *g*, 5 min), resuspended in 2% FBS/PBS, and used for flow cytometry or cell sorting.

#### Clodronate monocyte depletion

To deplete circulating monocytes, clodronate (clodronateliposomes.org, Vrije Universiteit, Netherlands) was injected *i.p.* into 12 w C57BL/6J mice daily for two days, while vehicle (PBS) control liposomes were used as control.

#### Hematopoietic colony-forming unit (CFU) assays

Clonogenicity was assessed by performing CFU cultures. Briefly, 2×10^5^ freshly isolated cells from digests of aorta or specified tissue were plated in methylcellulose (MethoCult GF M3434, StemCell Technologies), in duplicate. Where specified, cultures were performed in the presence or absence of Fractalkine (CX_3_CL1) (100 nM; Sigma-Aldrich), M-CSF (50 nM; PeproTech), AngII (10 nM, 100 nM, 1000 nM; Sigma-Aldrich), Enalapril (100 nM; Sigma-Aldrich), Losartan (100 nM; Sigma-Aldrich) or PD123,319 (50 nM; Sigma-Aldrich), which were replenished every three days. After 14 d, CFUs (defined as a minimum of 30 cells) were counted under light microscopy and classified by colony subtype (e.g. CFU-M) and size (in a single focal plane), as described previously ^34, 35^. In some experiments, after completion of CFU counts on day 14, colonies were isolated and disaggregated into single-cell suspensions for CFU-M renewal assays, flow cytometric characterization or other experiments as specified.

CFU renewal assays were performed by individually isolating 1° or 2° CFU-M from methylcellulose under light microscopy. Isolated colonies then disaggregated, and single-cells re-plated in methylcellulose in 96-well plates. Wells were imaged daily until day 14 to document and quantify the emergence of new colonies in 2° and 3° cultures from each plated cell.

CFU-M were also passaged in the presence or absence of 100 nM AngII in bulk culture. For this, the methylcellulose from 1° cultures was liquefied and disaggregated to obtain a single-cell suspension, which was then re-cultured in methylcellulose at a density of 2×10^5^ per well of a 24-well plate, for another 14 d before colony counting. Further passages (tertiary, quaternary, quinary) were conducted in the same way.

#### Flow cytometry and cell sorting

Single-cell suspensions from tissue digests or culture-derived CFU-M were resuspended in aliquots of ≤ 10 x 10^6^ cells/mL in PBS/5% FBS/0.2% sodium azide. 1 μL of fixable viability stain 700 (BD Horizon) was added to each sample and incubated for 15 min on ice. After washing, samples were blocked with purified rat anti-mouse CD16/CD32 (BD Pharmingen, San Diego CA, USA) antibody for 5 min on ice. Cells were then incubated for 30 min with 1⁰ antibodies and subsequently with 2⁰ antibodies when required (**Supplementary Table 15**). Samples were then washed and fixed in formalin/PBS for analysis with BD LSRFortessa^TM^ X-20 and BD FACs Diva Software (BD Biosciences) or Cytek^®^ Aurora and SpectroFlo^®^ software (Cytek Biosciences). For cell cycle analysis, BrdU Flow Kit (BD Biosciences) was used as per manufacturer’s instructions.

Cell populations in BM were defined by cell surface phenotypes as previously described ^41^. LT-HSCs were defined as Lin^−^Sca1^+^cKit^+^CD48^−^CD150/Slamf1^+^CD135/Flk2^−^, ST-HSCs^F^ as Lin^−^Sca1^+^cKit^+^CD135/Flk2^intermediate^ and MPPs as Lin^−^Sca1^+^cKit^+^CD48^+^CD150/Slamf1^−^ CD135/Flk2^+^.

Data files were analyzed using FlowJO version X software (Tree Star Inc., Ashland, OR, USA). Gating was performed based on SSC-A vs FSC-A (to exclude cell debris), FSC-H vs FSC-A (for single cells) and the use of Fixable Viability Stain 700 (BD Biosciences). We used FMO controls for each experiment to determine the positive percentage expression of different surface markers. A FACSAria Fusion cell sorter (BD Bioscience) was used for cell sorting.

#### Matrigel^TM^ cord-forming assay

FACS-sorted progenitors, macrophages and endothelial cells from 12 w C57BL/6J aortas, and progenitors isolated from day 14 medium-sized CFU-M were suspended in Endothelial Cell Growth Medium (Lonza, Cat# CC-4542) and plated on growth factor-reduced Matrigel^TM^ (BD Biosciences) at 2×10^4^ cells per well of a 96-well plate, to study their intrinsic capacity to form vascular-like cords. Cultures were photographed under a light microscope daily until day 7 to detect the presence of cords, identified as cellular extensions linking cell masses or branch points. Images were captured at 10x magnification in five different regions spanning the entire well. Total cord length and number of branching points were quantified using Image J software (NIH). Single-cell suspensions were retrieved from the Matrigel^TM^ following digestion with type IV collagenase (Sigma-Aldrich) for 45 min, neutralized with IMDM supplemented with 10% FBS, and immunostained for flow cytometry, as published ^53^.

For the single-cell differentiation assay, progenitors isolated from day 14 aortic CFU-M from UBI-GFP mice were disaggregated and serially diluted in Endothelial Cell Growth Medium to obtain a single GFP^+^ cell. A single GFP^+^ cell was plated on 15 µL of Matrigel^TM^ with 2×10^4^ GFP^−^ CFU-M-derived progenitors from C57BL/6J aortas per well in a µ-slide angiogenesis (ibidi^®^, cells in focus) and cultured for 7 d. Cultures were imaged on a Leica TCS SP8X/MP laser scanning confocal microscope (Leica Microsystems) on day 0 to demonstrate the presence of a single GFP^+^ cell in each well, and then on day 7 to determine its differentiation. Single-cell suspensions were prepared from Matrigel^TM^ cultures as described above. 100 µL of cell suspension from each well was placed in a Shandon single cytofunnel (EPKM5991040, Epredia^TM^) and centrifuged onto coated cytoslides (EPKM5991056, Epredia^TM^) in a Shandon Cytospin®4 Cytocentrifuge (Life Technologies, CA, US) (1000*g*, 5 min) for cytospin preparations for immunolabeling.

#### Aortic ring outgrowth model

Aortic ring assays were used to study the involvement of aortic EndoMac progenitor cells in adventitial angiogenesis. Aortic explants were carefully flushed to remove blood and dissected free of surrounding adipose. For studies from *Cx_3_cr1*^CreER-YFP^ x *Rosa*^tdTom^ mice which required intact adventitia, aortas were used with all three mural layers intact. For studies that required addition of GFP^+^ aortic EndoMac progenitors, C57BL/6J aortas were dissected to completely remove the adventitia. Rings of 1 mm thickness were then cut from ascending thoracic aorta, embedded in Matrigel^TM^ and overlaid with Endothelial Cell Growth Medium. Aortic sprouts were imaged on day 5 for quantification of adventitial sprout length using Image J software. Single-cell suspensions were retrieved from adventitial sprouts in Matrigel^TM^ as described above, and immunostained for flow cytometry.

#### Wright-Giemsa staining

FACS-sorted progenitors, macrophages and endothelial cells from aortas of E9.5 TAM-induced 12 w *Cx_3_cr1*^CreER-YFP^ x *Rosa*^tdTom^ mice or E8.5 TAM-induced 12 w *Csf1r*^Mer-iCre-Mer^ x *Rosa*^mTmG^ mice were resuspended in 200 μL of IMDM containing 2% FBS. Cells were cytospun (300 *g*, 5 min) onto cytoslides using a cytospin centrifuge (Thermo Fisher) and then stained with Wright-Giemsa reagent (Sigma-Aldrich) followed by imaging on a light microscope (Carl Zeiss, Axio).

#### Immunofluorescence labeling and confocal microscopy

Intact tissue samples were harvested from mice and placed in 30% sucrose overnight, fixed with 10% formalin for 24 h then embedded in Optimal Cutting Temperature (O.C.T.) compound (Sakura Finetek). Five µm thick frozen sections were cut, fixed and then placed in methanol with 0.3% H_2_O_2_ before heat-mediated citrate antigen retrieval. Blocking was performed with either 10% normal goat or donkey serum, or 3% normal horse serum. Sections were incubated with 1° antibodies overnight followed by 2° antibodies (**Supplementary Table 15**). Nuclei were counterstained with DAPI. Microscopy was performed with a Leica TCS SP8X/MP laser scanning confocal microscope (Leica Microsystems).

For labeling of cytospun preparations, slides were fixed in 4% paraformaldehyde (PFA) for 10 min. Permeabilization was performed with 0.1% Triton X-100 and blocking with 5% normal horse serum (NHS) for 30 min each, and then hybridisation with 1⁰ antibody overnight at 4⁰C followed by 2⁰ antibody for 45 min at 37⁰C. All antibodies were diluted in 1% NHS. Nuclei were counterstained with DAPI. Confocal microscopy was performed as above. Images were taken from 20-25 fields of view to image all cells and analyzed using ImageJ software.

#### Hindlimb ischemia model

For adoptive cell transfer studies, we used male and female 12 w C57BL/6J mice as recipients and UBI-GFP mice as donors. To induce hindlimb ischemia, the proximal portion of the left femoral artery and the distal saphenous artery were both ligated, as well as the popliteal artery and all branches in between, and an arterectomy performed to remove the intermediate segment of vessel. Purity checked EndoMac progenitor cells isolated from medium CFU-M grown from either adult aortic cells or E9.5 YS cells from GFP donor mice (7.5×10^3^ in 60 μL Matrigel^TM^) were administered as three intramuscular injections into the quadriceps and gastrocnemius of the ischemic limb. A control group received injections of cell-free Matrigel^TM^.

Hindlimb blood flow reperfusion was determined by laser Doppler perfusion imaging (moorLD12-IR, Moor Instruments) immediately following surgery and then at days 7 and 14 post-ischemic induction. Euthanasia was performed after the final imaging on day 14. Peripheral blood was taken by cardiac puncture, while the quadriceps and gastrocnemius muscle from both hindlimbs was harvested and processed for hematopoietic CFU assays, flow cytometry or immunofluorescence staining, to detect engraftment of donor GFP^+^ cells, their fate and vasculogenic capacity.

#### Single cell RNA sequencing and data analysis

Single cell RNA sequencing (scRNA-seq) was performed on viable (DAPI^+^) aortic CFU-M progenitors grown from two 12 w C57BL/6J male mice. Isolated progenitors were subjected to MULTI-seq labeling as previously described ^76^. Briefly, cells were resuspended in 18 µL of 1X PBS, labeled with a unique 2 µM anchor lipid-modified oligonucleotide (LMO)-Barcoded Oligo complex for 5 min on ice. After this incubation, 2 µM co-anchor was added and cells incubated on ice for another 5 min. Labeling was quenched with 1 mL of PBS + 2% BSA and cells from each sample in the run mixed for washing the pool with PBS + 1% BSA (twice); final resuspension was in PBS + 1% BSA. From pooled cells, single cell sequencing libraries were prepared using the Chromium Next GEM single-cell 3’ kit (v3.1, 10x Genomics) and Chromium system (10x Genomics) according to manufacturer’s specifications, at the South Australian Genomics Centre (SAGC, SAHMRI, Adelaide). Paired-end sequencing was performed first, on an Illumina Nextseq High using v2 chemistry, at the SAGC, and subsequently, to increase read depth, on an Illumina Novaseq 6000 (Illumina, San Diego, USA) using one lane of an S4 flowcell (Illumina), at the Australian Genome Research Facility (AGRF, Melbourne, Australia).

Cell Ranger v7.0.0 (10x Genomics) was used to process raw sequencing data from the two runs before subsequent analyses. This pipeline aligned sequencing reads from FASTQ files to the mm10 transcriptome using the STAR aligner and quantified the expression of genes in each cell. FASTQ files from the two sequencing runs were then combined and cell ranger analysis performed on combined FASTQ files to quantify gene expression and barcode enrichments per cell. The Seurat demultiplexing pipeline ^77^ was applied to demultiplex cells to their original sample-of-origin. Cells detected with zero barcodes (Negatives) or more than one barcode (Doublets) were discarded in this analysis. From the remaining 16,350 singlets, cells containing expression values for <200 genes or >9000 genes were filtered out as initial quality control measures. Moreover, cells with more than 20% of mitochondrial genes were filtered out to exclude low-quality and dying cells, which often exhibit extensive mitochondrial contamination. These resulted in 16,088 singlets corresponding to three barcodes, BC1: aortic CFU-M population (7,977 cells), BC2: Study population 2 (4,795 cells), and BC3: Study population 3 (3,316 cells). The data from the latter two populations will be published elsewhere. A further quality control measure of 10% mitochondrial gene cut-off was applied to the aortic CFU-M population. The remaining cells were then used to characterize the cellular heterogeneity of CFU-M progenitors. Processed scRNA-seq data was analyzed in R v4.1.3 using the Seurat R package v4.1.1 ^77^.

Dimensionality reduction was performed using Principal Component Analysis to explore transcriptional heterogeneity and perform clustering of cell types. Twenty principal components (PCs) were selected based on the elbow plot. PC loadings were used as inputs for a graph-based approach to cluster cells by cell type with resolution 0.6, and as inputs for Uniform Manifold Approximation and Projection (UMAP) for visualization purposes. Marker genes for each cluster were identified using “FindAllMarkers” R function. Phylogenetic tree relating the ‘average’ cell from each CFU-M identity class was constructed using “BuildClusterTree” R function. ggtree R package ^78^ was then used for visualization and annotation of the CFU-M cluster tree. Trajectory analysis was performed using Monocle 3^79–81^ as described in http://cole-trapnell-lab.github.io/monocle3/. Monocle 3 R package offers trajectory analysis to model the relationships between groups of cells as a trajectory of gene expression changes. The pseudotime, a measure of progression over trajectory, was calculated with CFU-M trajectory starting from Progenitors1. The graph test function of Monocle was utilised to identify genes that vary over the CFU-M trajectory. Figures were primarily generated using Seurat, ggplot2 ^82^ and ggtree R packages.

Bioinformatic analysis of marker genes for each cluster was performed using Innate DB. To identify pathways and transcription factor binding sites associated with marker genes, pathway, and transcription factor binding site overrepresentation analyses were performed using the hypergeometric algorithm and Benjaminin & Hochberg correction for multiple testing with FDR of 0.05.

#### Gene expression analysis by RT-PCR

Total RNA was extracted from 12 w C57BL/6J aortic digests or aortic CFU-M progenitors, including after exposure to AngII, using Total RNA kit (Bio-Rad). 200 ng of total RNA was reverse transcribed using iScript cDNA synthesis kit (Bio-Rad). Quantitative real-time PCR was performed using the SsoAdvanced™ Universal SYBR® Green RT-PCR Kit (Bio-Rad) on a CFX96 thermocycler (Bio-Rad). Relative changes in target gene expression were normalized using the ^ΔΔ^Ct method to the housekeeping gene *Actb* (β-actin) and the experimental control condition. Primer sequences used in this study are listed in **Supplementary Table 16**.

#### Peritoneal transfer assay

Progenitor cells were isolated from medium CFU-M grown from 12 w UBI-GFP aortas and checked for purity. 1 x 10^4^ GFP^+^ progenitors were injected into the peritoneal cavity of 12 w C57BL/6J mice, which were then randomised to receive daily *i.p.* injections of either PBS, 0.7 mg/kg AngII or 0.05 μg M-CSF. Mice were euthanized after 72 h and cells harvested from the peritoneal cavity by injecting 5 mL of PBS *i.p..* Following gentle massage of the peritoneum, the peritoneal fluid was collected and centrifuged (400 *g*, 5 min) to obtain single-cell suspensions for immunostaining and analysis of GFP^+^ cells by flow cytometry.

#### Angiotensin II infusion studies

Osmotic minipumps containing AngII (0.7 mg/kg/d) or equivalent volume of PBS, were implanted subcutaneously between the shoulder blades of E9.5 TAM-induced adult *Cx3cr1*^CreER-YFP^ x *Rosa*^tdTom^ mice or adult *Flt3*^Cre^ x *Rosa*^mTmG^ mice, randomly selected for treatment. Cohorts of mice were euthanized at specified time-points.

#### Statistical analysis

Where relevant, data analysis was performed blinded to study group. Statistical analyses were performed using Prism (GraphPad). Data sets were tested for normality of distribution by Shapiro-Wilk test. The ROUT method was applied to identify and exclude any outliers, with Q set to 1%. Statistical comparisons were performed with parametric or non-parametric unpaired or paired two-sample *t*-tests or ANOVA (with post-test multiple comparisons), as specified. Results are expressed as mean ± standard deviation of multiple experiments. In all cases, statistical significance was established at two-tailed p < 0.05. The statistical parameters and the number of mice used per experiment are given in the figure legends.

#### Data availability

The single-cell RNA-seq data has been deposited in the National Centre for Biotechnology Information Gene Expression Omnibus ^83^ and is accessible through GEO Series accession number GSE232625 (https://www.ncbi.nlm.nih.gov/geo/query/acc.cgi?&acc=GSE232625). All other data will be made available upon request.

## ACKNOWLEDGEMENTS

The authors thank Adelaide Microscopy at the University of Adelaide and staff of the Bioresources Facility, Flow Cytometry Core Laboratory and South Australian Genomics Centre of the South Australian Health and Medical Research Institute, especially Mr Randall Grose and Mr Mark Van der Hoek.

This work was supported by the National Health and Medical Research Council of Australia [PG 1086796, IG 2001541, PRF 1111630 to S.J.N., CDF 1161506 to P.J.P.]; the National Heart Foundation of Australia [VG 102981, Lin Huddleston Fellowship to C.A.B., FLF 100412, 102056 and 106656 to P.J.P.]; and the Royal Australasian College of Physicians.

## AUTHOR CONTRIBUTIONS

Conceptualization, A.W. and P.J.P.; Investigation, A.W., S.L., M.H., M.I.D., D.T-F., D.X.A.T., C.D., N. S., S.F., T.S., A.L., B.A.D.B., J.K., N.L.H., G.R.D., A.V., V.C., A.M., Z.N., J.T.M.T., L.M., A.R.P., S.S. and P.J.P.; Writing – Original draft, A.W. and P.J.P.; Writing – Review & Editing, A.W., N.L.H., G.R.D., V.C., J.T.M.T., J.M.P., C.S.B., A.R.P., S.S., S.J.N, C.A.B. and P.J.P.; Supervision, C.S.B., A.R.P., C.A.B. and P.J.P.; Funding acquisition, P.J.P..

## EXTENDED DATA FIGURES

**Extended Data Fig. 1.**
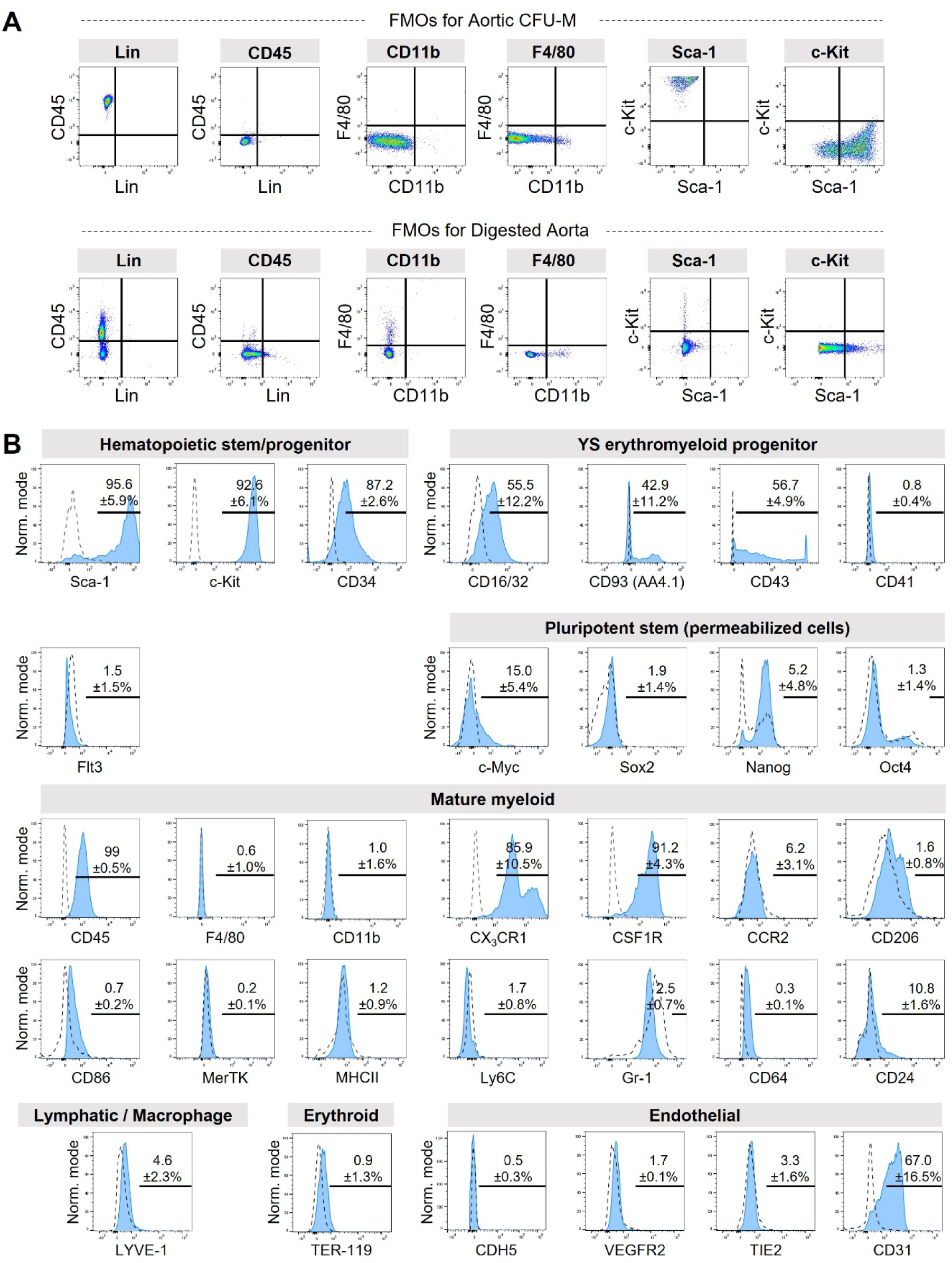
Surface marker expression of progenitors contained within 1° aortic CFU-M. (A) Top row: Fluorescence-minus-one (FMO) controls for flow cytometry performed on cells from medium-sized aortic CFU-M in **Fig. 1C** and **D**. Bottom row: FMO controls used for flow cytometry performed on cells from aortic digests in **Fig. 1E**. (B) Flow cytometry histograms show expression of different surface markers on cells from medium-sized 1° aortic CFU-M (blue histogram). Dotted histogram, FMO control. Mean±SD % expression values are shown from n≥3 12 w C57BL/6J mice. Also see **Fig. 1**.

**Extended Data Fig. 2.**
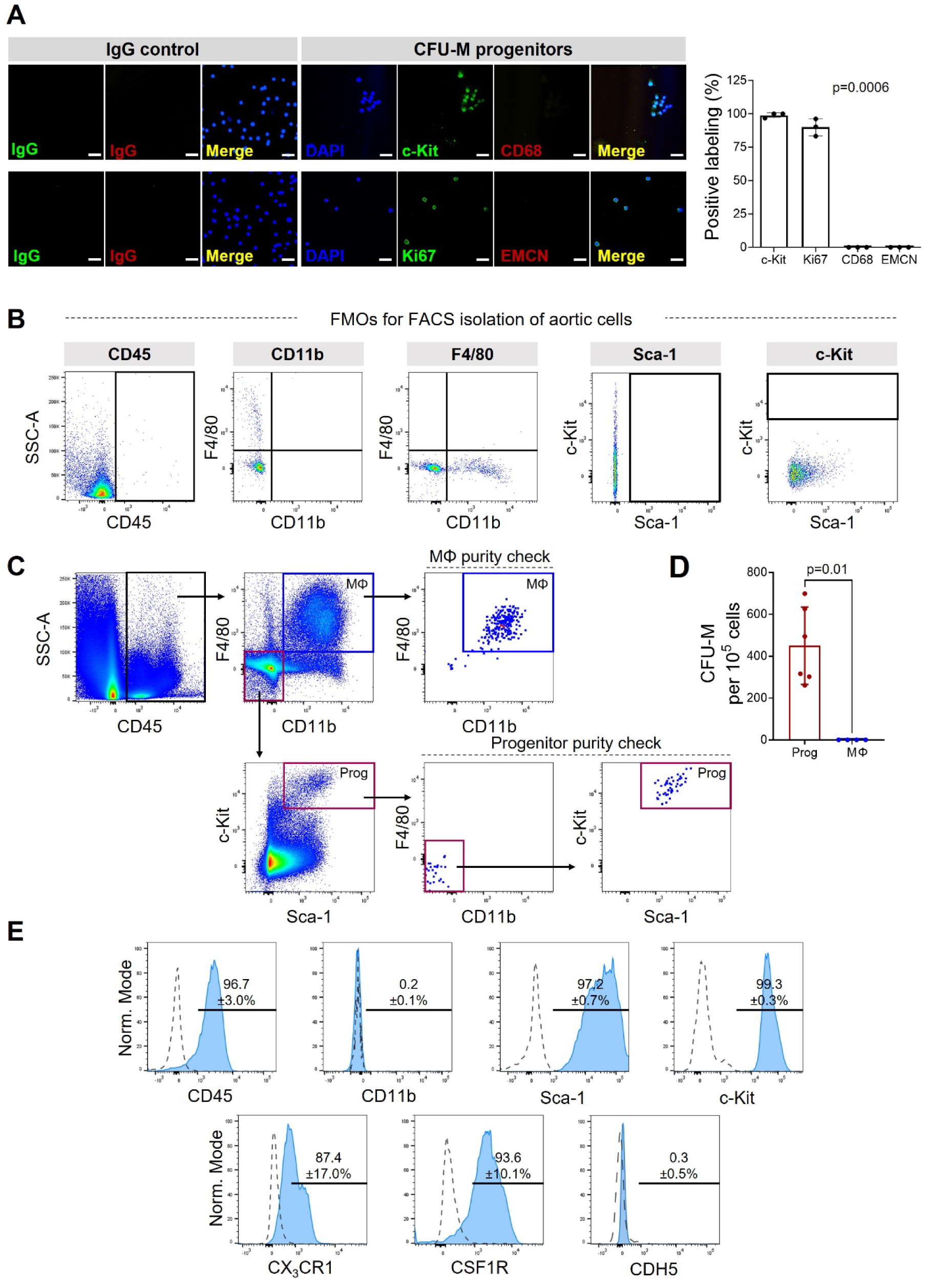
Immunophenotypic confirmation of cell population responsible for aortic CFU-M. (A) Confocal microscopy images of immunofluorescence labeling of cytospin preparations of cells from aortic CFU-M. Scale bar, 100µm. Graph summarizes % of positively labeled cells with each marker. Data from three separate experiments using CFU-M from adult C57BL/6J aorta. Kruskal-Wallis test. (B) FMO controls used for FACS to isolate progenitors (Prog) and macrophages (MΦ) from aortic cells of adult C57BL/6J mice. (C) Representative flow cytometry dot plots used to FACS isolate progenitors and macrophages from aorta, with purity checks of sorted populations. (D) Graph summarizes CFU-M yield from sorted aortic cell populations. N=4-6 independent experiments each using aortas from ≥6 mice. Mann-Whitney test. (E) Flow cytometry histograms showing surface marker expression of cells contained in CFU-M produced by sorted aortic progenitors (blue). Dotted histograms, FMO controls. N=3. Data summarized as mean±SD. Also see **Fig. 1**.

**Extended Data Fig. 3.**
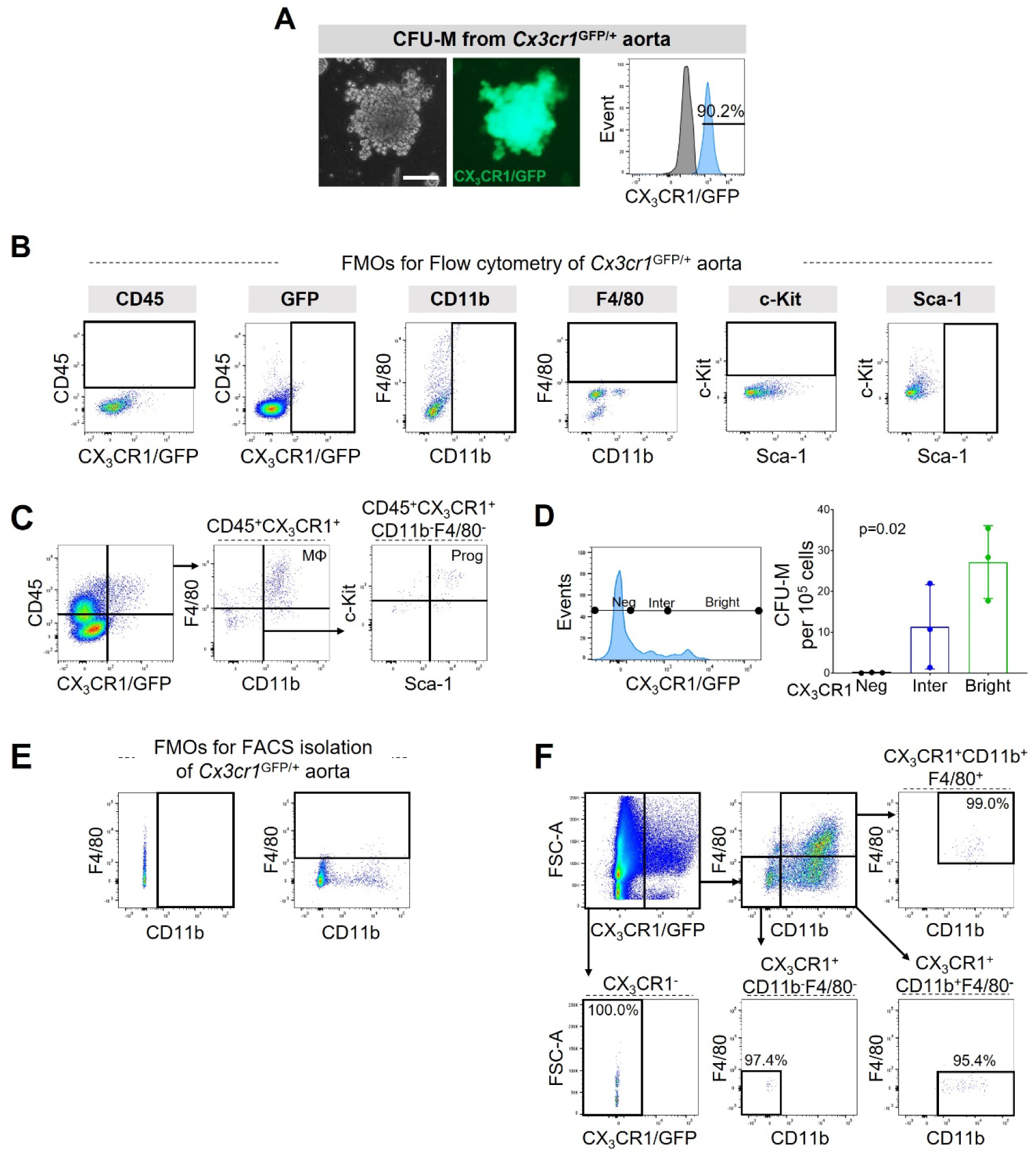
Aortic CFU-M progenitors express CX_3_CR1. (A) Light and fluorescence microscopy and flow cytometry (blue histogram) show GFP^+^ (CX_3_CR1^+^) expression of cells contained in CFU-M from 12 w *Cx3cr1*^GFP/+^ aortas. Gray histogram, C57BL/6J control. N=1. Scale bar, 100 µm. (B) FMO controls used for flow cytometry analysis of aortic digests from *Cx3cr1*^GFP/+^ mice. (C) Flow cytometry shows presence of CD11b^−^F4/80^−^Sca-1^+^c-Kit^+^ progenitors and CD11b^+^F4/80^+^ macrophages among CD45^+^CX_3_CR1^+^ cells in aortas of 12 w *Cx3cr1*^GFP/+^ mice. Representative of n=6. (D) FACS isolation of CX_3_CR1^Neg^, CX_3_CR1^Inter^ and CX_3_CR1^Bright^ populations from 12 w *Cx3cr1*^GFP/+^ aortas. Graph shows 1° CFU-M yield from each population (three experiments, six aortas each). Repeated measures one-way ANOVA. (E) FMO controls used for FACS isolation of aortic cells from *Cx3cr1*^GFP/+^ mice. (F) Gating strategy and purity checks for FACS isolation of aortic cells from 12 w *Cx3cr1*^GFP/+^ mice into CX_3_CR1^+^CD11b^−^F4/80^−^, CX_3_CR1^+^CD11b^+^F4/80^+^, CX_3_CR1^+^CD11b^+^F4/80^−^ and CX_3_CR1^−^ populations, used in **Fig. 1H**. Data summarized as mean±SD. Also see **Fig. 1**.

**Extended Data Fig. 4.**
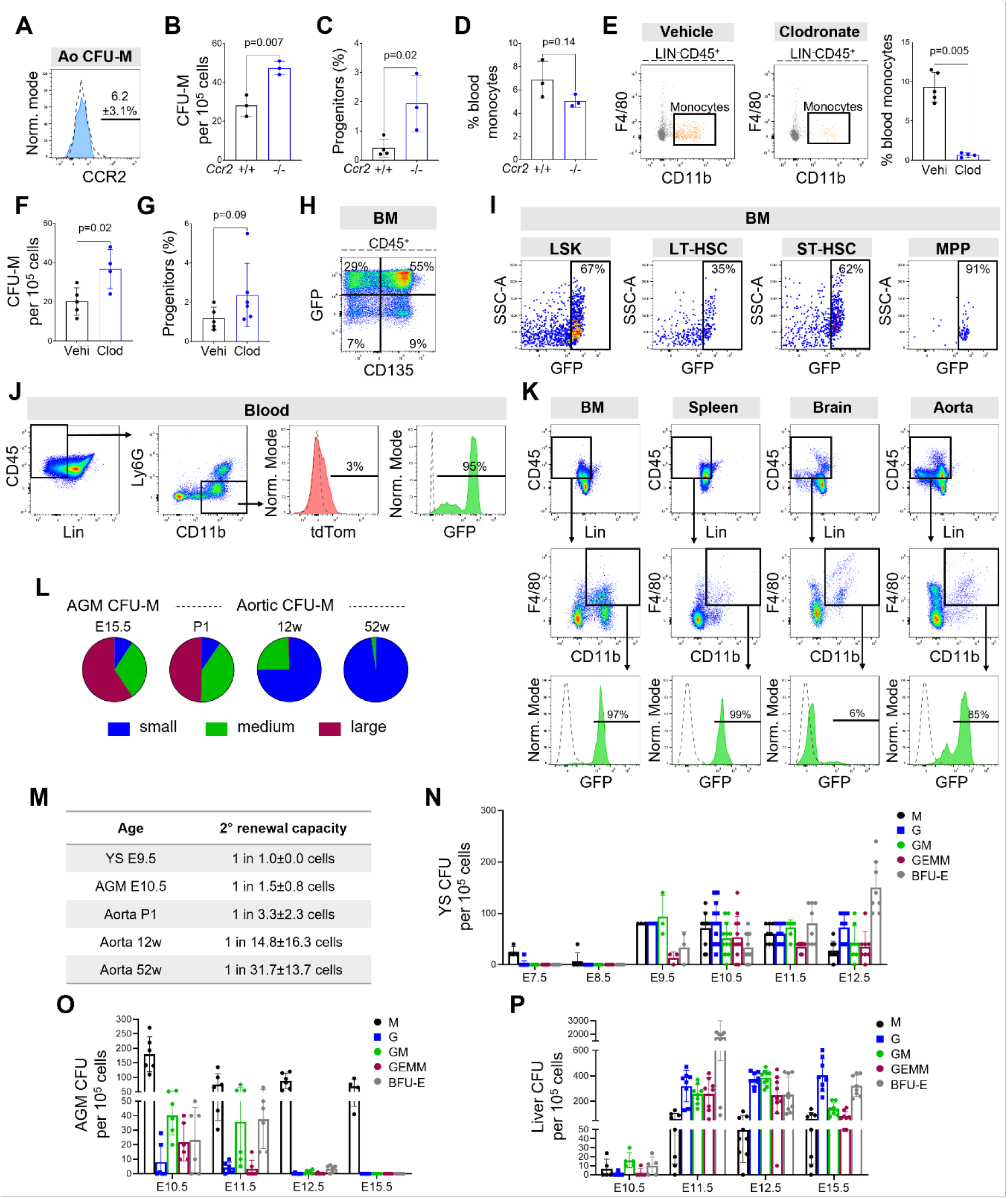
Origins of CFU-M. (A) Surface expression of CCR2 on progenitor cells from 1° aortic CFU-M (blue histogram). Dotted histogram, FMO control. N=3. (B-D) Graphs show comparisons of (B) CFU-M yield and (C) percentage of progenitors in aorta, and (D) blood monocytes between 12 w *Ccr2*^+/+^ and *Ccr2*^−/−^ mice, measured by flow cytometry (n=3; unpaired t-tests). (E-G) Flow cytometry plots and graphs showing comparisons of (E) percentage of blood monocytes, and (F) CFU-M yield and (G) percentage of progenitors in aorta of 12 w C57BL/6J mice following treatment with clodronate (Clod) or vehicle (Vehi) (n=4-6). Unpaired t-tests for (E) and (F) and Mann-Whitney test for (G). (H) Flow cytometry plot showing correlation of CD135 (Flt3) and GFP expression in CD45^+^ cells in BM of adult *Flt3*^Cre^ x *Rosa*^mT/mG^ mice (n=5). (I-J) Flow cytometry plots and histograms show GFP^+^ (Flt3-cre^+^) status of (I) BM long-term (LT) and short-term (ST) HSCs, multipotent progenitors (MPP) and (J) blood monocytes from adult *Flt3*^Cre^ x *Rosa*^mT/mG^ mice. Percentages represent mean of n=5-6. Dotted histograms, C57BL/6J control. LSK, Lin^−^Sca-1^+^c-Kit^−^. *Please see Source Data file for all FMO controls and gating strategy for BM, and Methods for immunophenotypic definitions of BM cell populations*. (K) Flow cytometry of GFP^+^ (Flt3-Cre^+^) expression on macrophages from different tissues of adult *Flt3*^Cre^ x *Rosa*^mTmG^ mice. Green histogram, sample. Dotted histogram, control. Percentages shown are mean % GFP^+^ cells from n=3-4 mice. (L) Pie charts show classification of CFU-M based on colony size from E15.5 AGM and aorta at different postnatal ages from C57BL/6J mice. N=3-6 mice per age. (M) Frequency of self-renewal of progenitor cells from 1° CFU-M from YS, AGM and aorta at different ages, expressed as the number of individually seeded cells required to generate a 2° CFU-M. N=3-20 replicates per age. (N-P) Frequency of different CFU types from (N) YS, (O) AGM and (P) liver at different embryonic ages. N=1-6 C57BL/6J embryos per gestational age. M, macrophage; GM, granulocyte-macrophage; GEMM, granulocyte-erythrocyte-monocyte-megakaryocyte; G, granulocyte; BFU-E, burst-forming units-erythroid. Data summarized as mean or mean±SD. Also see **Fig. 2**.

**Extended Data Fig. 5.**
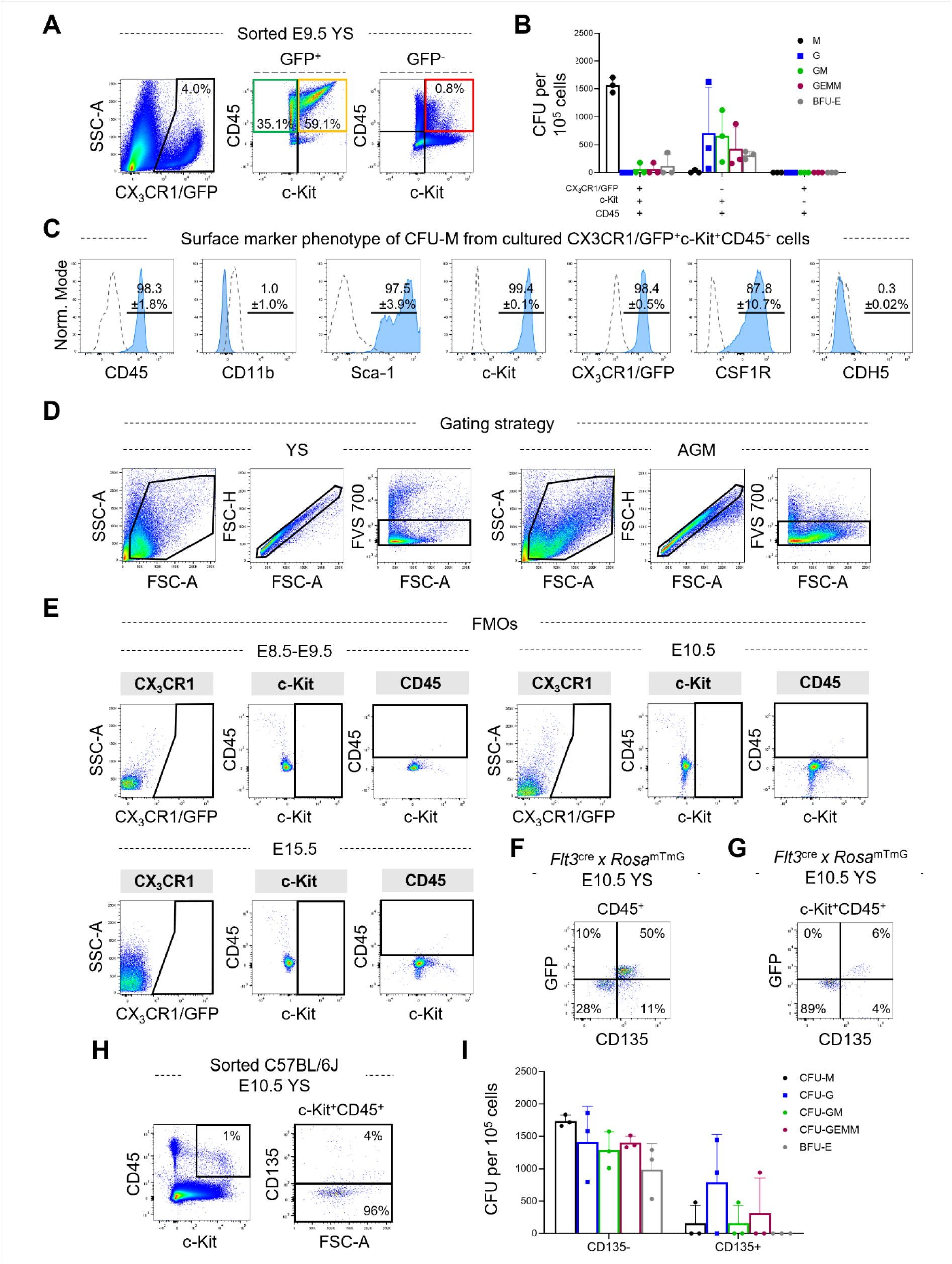
Identification of CFU-M progenitors in E9.5 Yolk Sac. (A) Gating strategy used for FACS isolation of YS cells from E9.5 *Cx3cr1*^GFP/+^ embryos into CX_3_CR1/GFP^+^c-Kit^+^CD45^+^ (yellow box), CX_3_CR1/GFP^+^c-Kit^−^CD45^+^ (green box) and CX_3_CR1/GFP^−^c-Kit^+^CD45^+^ (red box) populations, used in **Fig. 2G**. Percentages in quadrants are from the parent population and relate to the example shown. (B) Yield of different colony types from the FACS-isolated populations. N=3 experiments. Also see **Fig. 2G**. (C) Flow cytometry histograms showing surface expression of markers (blue histograms) on cells contained in CFU-M produced by sorted YS progenitors. Dotted histograms, FMO controls. N=2-4. (D) Gating strategy used for flow cytometry of YS and AGM cells from *Cx3cr1*^GFP/+^ embryos shown in **Fig. 2H**. (E) FMO controls used for flow cytometry and FACS isolation of YS and AGM cells from *Cx3cr1*^GFP/+^ embryos in **Fig. 2H** and **Extended Data Fig. 5**. (F-G) Flow cytometry plots showing correlation of CD135 (Flt3) and GFP expression in (F) CD45^+^ and (G) c-Kit^+^CD45^+^ cells in E10.5 YS from *Flt3*^Cre^ x *Rosa*^mT/mG^ embryos (n=5). (H) Gating strategy used for FACS isolation of c-Kit^+^CD45^+^CD135^−^ and c-Kit^+^CD45^+^CD135^+^ cells from E10.5 YS from C57BL/6J embryos. Percentages are from the parent population of the example shown. (I) Graph summarizes the yield of different types of colonies formed from the two FACS-isolated cell populations (n=3 pools each consisting of tissue from 4-10 embryos from >1 litter). Data summarized as mean or mean±SD. Also see **Fig. 2**.

**Extended Data Fig. 6.**
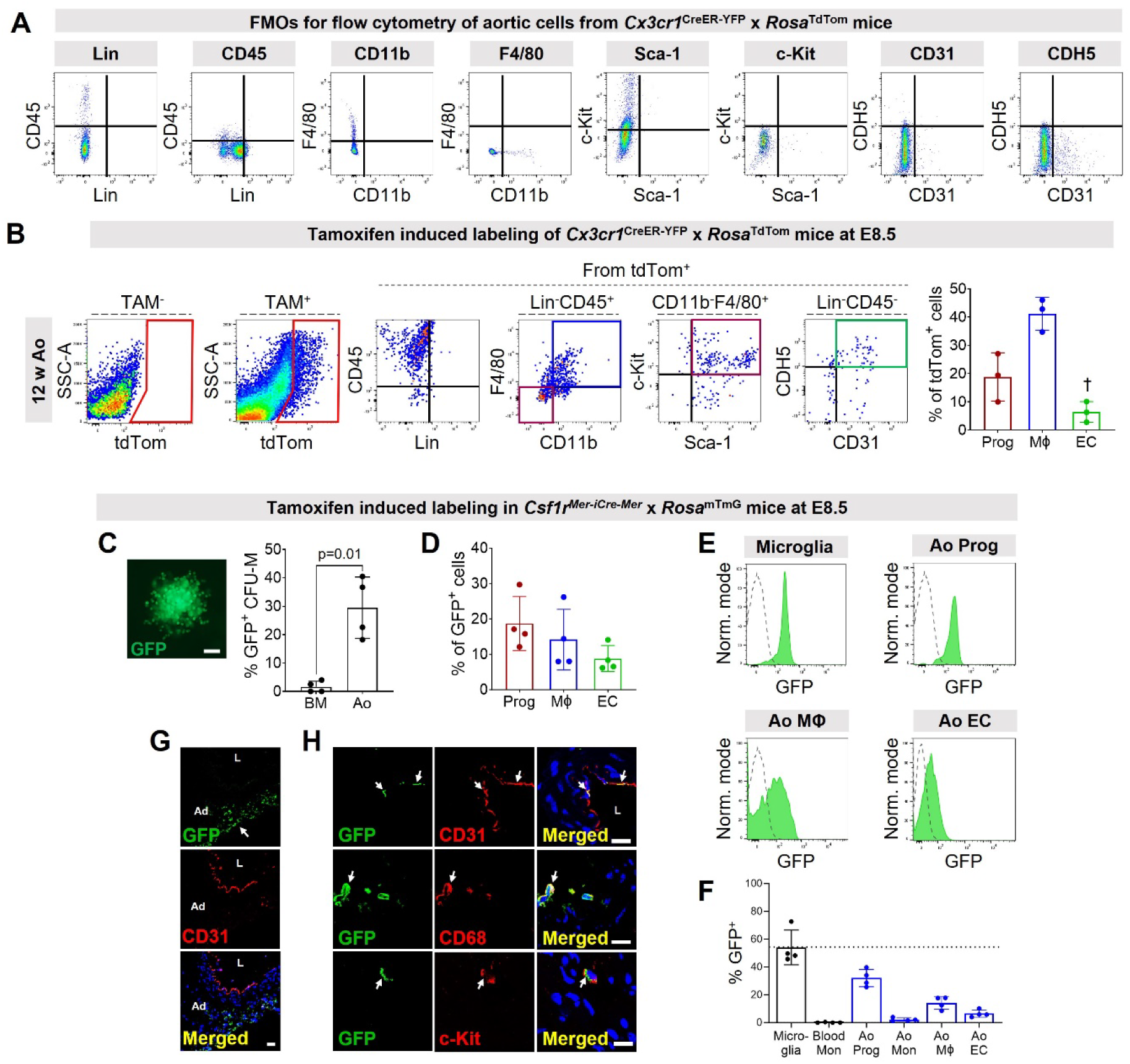
Aortic CFU-M progenitors arise from CX3CR1*+* and CSF1R*+* cells in the early embryo. (A) FMO control staining used to analyze cells from E15.5 AGM and adult aorta from *Cx3cr1*^CreER-YFP^ x *Rosa*^tdTom^ mice (also see **Fig. 3C,D**). (B) Flow cytometry dot plots show composition of tdTom^+^ cells in aorta of adult *Cx3cr1*^CreER-YFP^ x *Rosa*^tdTom^ mouse that had been induced with TAM *in utero* at E8.5. Gated regions correspond to macrophages (Mϕ, blue), progenitors (Prog, maroon) and endothelial cells (EC, green). tdTom expression from aorta of no tamoxifen (TAM^−^) control also shown. Graph shows the % of tdTom^+^ cells that comprised each of these three cell populations (n=3). Repeated measures ANOVA p=0.02. Multiple comparisons test: †p<0.01 vs macrophages. (C-H) *Csf1r*^Mer-iCre-Mer^ x *Rosa*^mTmG^ mice were induced with TAM at E8.5 and analyzed at 12 w. (C) Image of GFP^+^ CFU-M from 12 w aorta. Graph shows % GFP^+^ CFU-M from aorta and BM (n=4; paired t-test). (D) Graph summarizes % breakdown of GFP^+^ cells in adult aorta (n=4). Repeated measures ANOVA p=0.16. (E, F) Histograms and graph show % GFP expression in different cell populations from 12 w mice (n=4). Green histogram, sample. Dotted histogram, TAM^−^ control. (G, H) Confocal microscopy of immunolabeled sections of 12 w aorta shows (G) GFP^+^ cells in adventitia (Ad), with (H) magnified examples of GFP^+^ cells expressing CD31 in the intima and CD68 and c-Kit in adventitia (arrows). L, lumen. Data summarized as mean±SD. Scale bar, 100 μm in (A) and 20 μm in (G, H). Also see **Fig. 3**.

**Extended Data Fig. 7.**
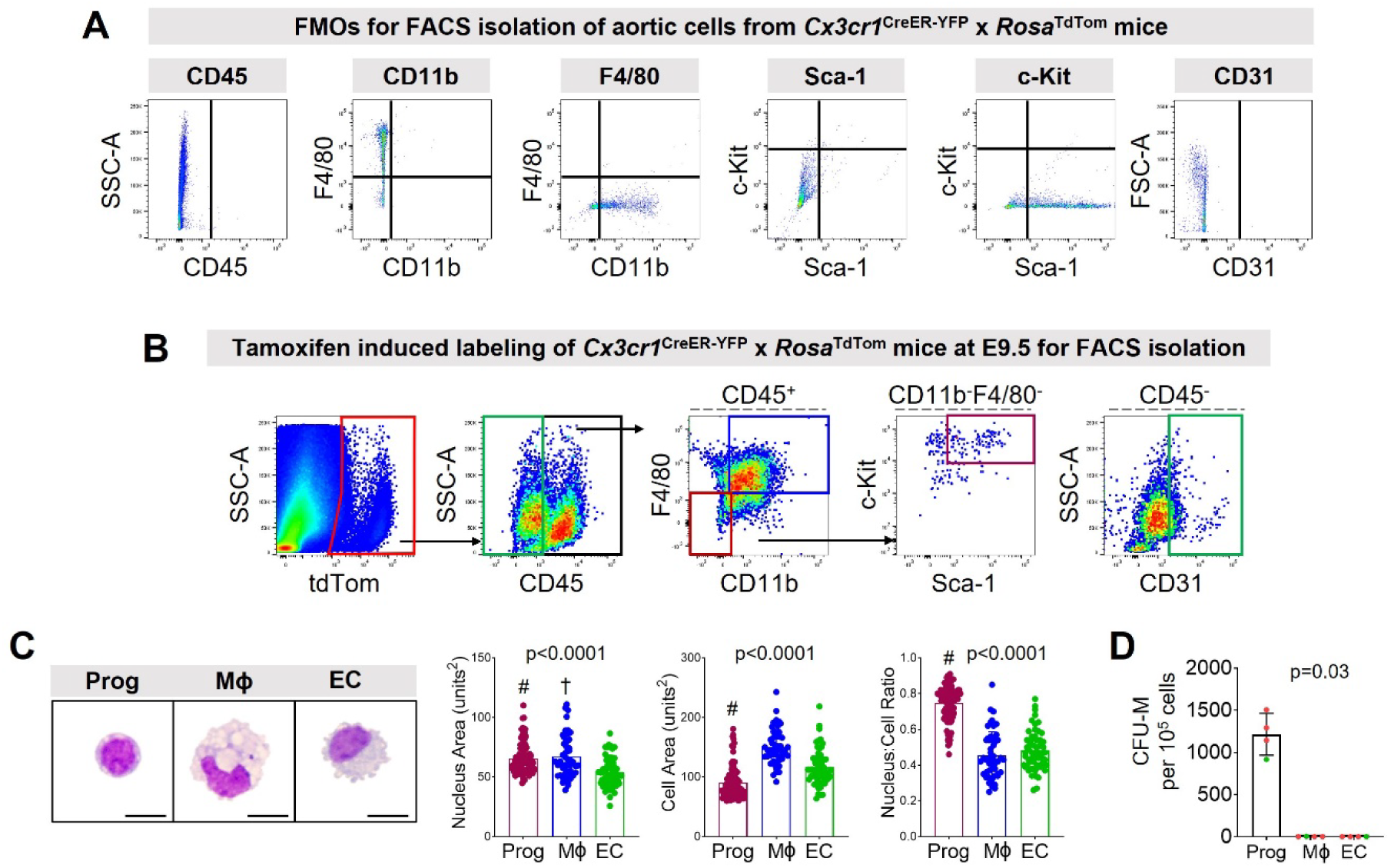
Embryonic CX_3_CR1^+^ cells give rise to EndoMac progenitors in postnatal aorta. (A-D) *Cx3cr1*^CreER-YFP^ x *Rosa*^tdTom^ mice were induced with TAM at E9.5 and used for FACS isolation of tdTom^+^ aortic cells at 12 w. (A) FMO controls used to sort aortic cells. (B) FACS isolation of tdTom^+^ progenitors, macrophages and endothelial cells from 12 w aorta. (C) Wright-Giemsa staining of sorted cell populations, and graphs showing comparisons of total cell area, nucleus area and nucleus:cell area ratio (n=51-100 cells per cell type). Kruskal-Wallis tests. Multiple test comparisons: #p<0.0001 vs Mϕ and EC; †p<0.01 vs EC. (D) CFU-M yield from tdTom^+^ (red dots) and GFP^+^ (green dots) progenitors, macrophages and endothelial cells isolated from adult aortas of E9.5 TAM-induced *Cx3cr1*^CreER-YFP^ x *Rosa*^tdTom^ mice or E8.5 TAM-induced *Csf1r*^MerCreMer^ x *Rosa*^mTmG^ mice, respectively (total of four experiments, using six aortas each). Friedman test. Data summarized as mean±SD. Also see **Fig. 3**.

**Extended Data Fig. 8.**
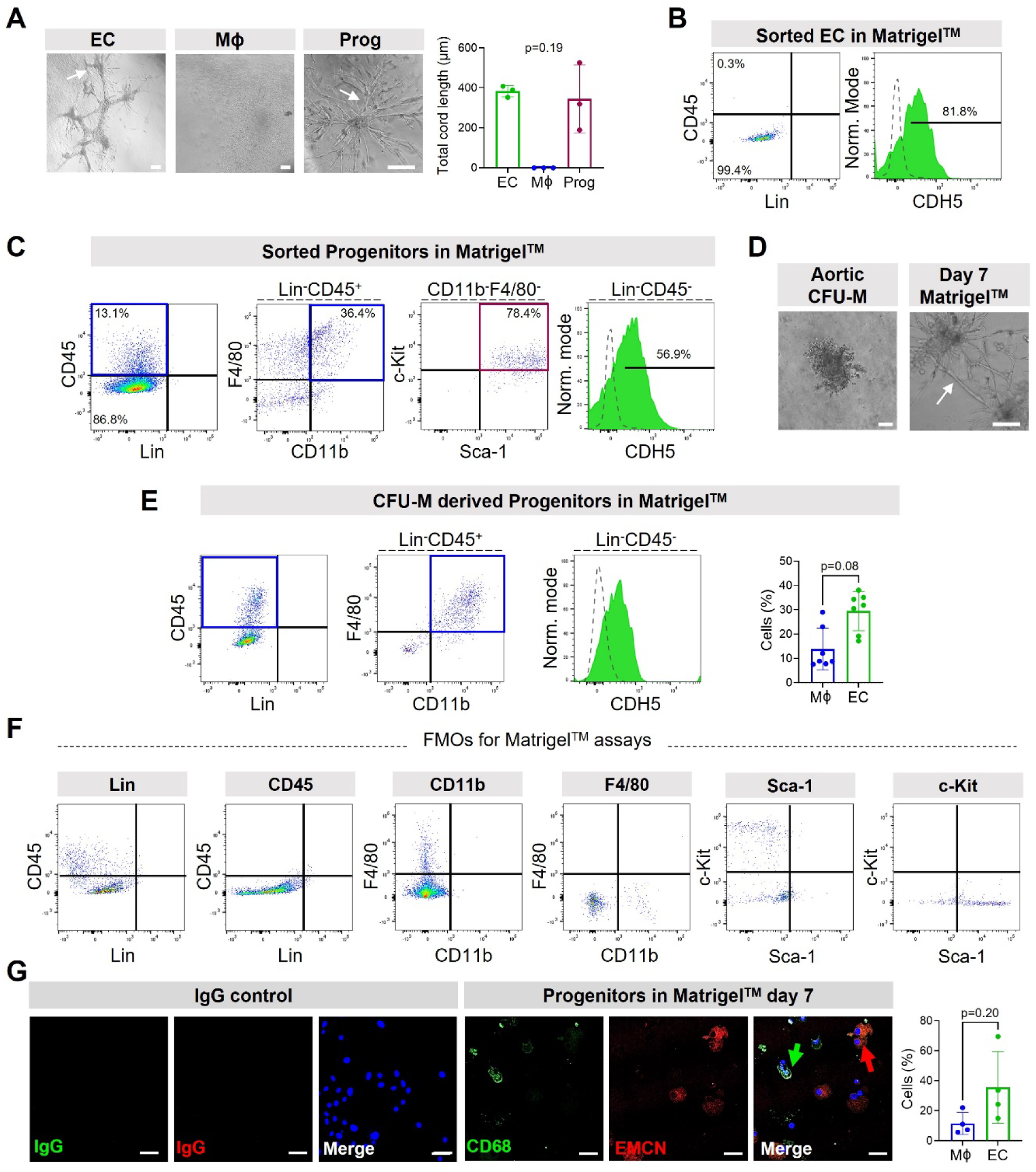
Aortic CFU-M progenitors have endothelial and macrophage differentiation potential. (A) Light microscopy images of FACS-isolated endothelial cells (EC), macrophages (Mϕ) or progenitors (Prog) from 12 w C57BL/6J aorta cultured in Matrigel^TM^ for 7 d. Arrows indicate cords. Graph shows total cord length (three experiments, six aortas each). Friedman test. (B, C) Flow cytometry plots and histograms show the cell types produced from culturing FACS-isolated primary aortic (B) endothelial cells and (C) progenitors in Matrigel^TM^ (n=1 experiment using six pooled aortas). Green histogram, sample; dotted histogram, FMO control. (D) Example of an aortic CFU-M and branching cord networks (arrow) produced from CFU-M derived progenitors in Matrigel^TM^. (E) Flow cytometry plots and graph show the cells produced from CFU-M progenitors cultured in Matrigel^TM^ for 7 d (n=7 adult C57BL/6J aortas, three independent experiments). Wilcoxon signed-rank test. (F) FMO controls for (B), (C) and (E) above. (G) Confocal microscopy of immunolabeled cytospin preparations of cells produced by CFU-M progenitors cultured in Matrigel^TM^ for 7 d. Cells stained with nuclear dye DAPI (blue) and immunolabeled for CD68 (green) and endomucin (EMCN; red). Graph summarises frequency of CD68^+^ and EMCN^+^ cells (n=4 independent experiments). Paired *t*-test. Data summarized as mean±SD. Scale bar, 40 µm in (A, D) and 100 µm in (G). Also see **Fig. 4**.

**Extended Data Fig. 9.**
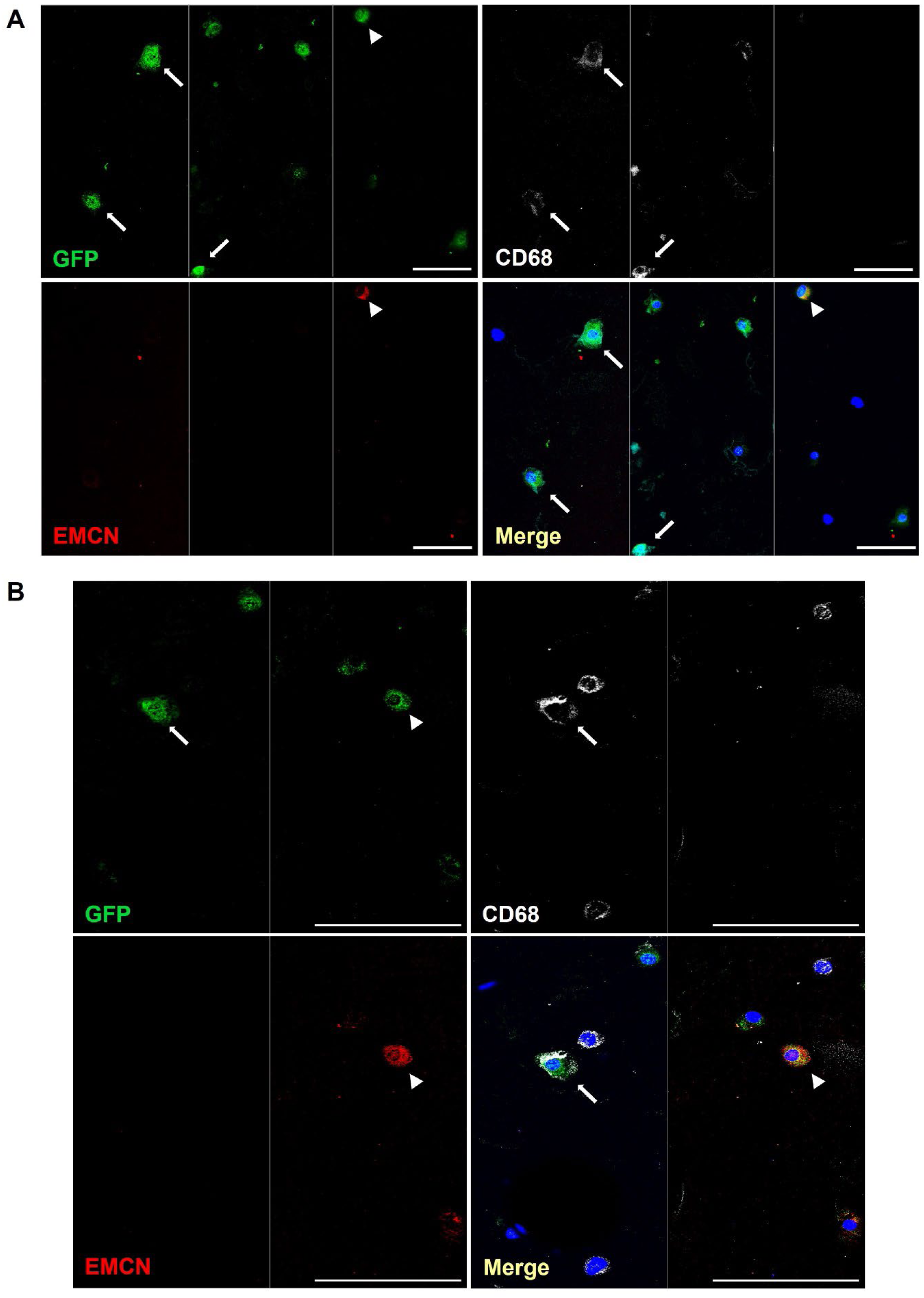
Aortic CFU-M progenitors are bipotent for endothelial cells and macrophages at a single cell level. (A, B) A single GFP^+^ aortic CFU-M progenitor cell was seeded with GFP^−^ progenitors on Matrigel^TM^ in individual wells and cultured for 7 d. Cytospun preparations of resulting sprouts were immunolabeled as indicated. Confocal microscopy images of immunolabeled cells from two independent wells (A and B) show presence of GFP^+^CD68^+^ macrophages (arrows) and GFP^+^EMCN^+^ endothelial cells (arrowheads). GFP^+^CD68^−^EMCN^−^ cells and GFP^−^ CD68^+^ macrophages can also be seen. A montage of 2-3 fields of view per well is presented. White vertical lines demarcate the fields of view. Scale bar, 100 µm. Also see **Fig. 4**.

**Extended Data Fig. 10.**
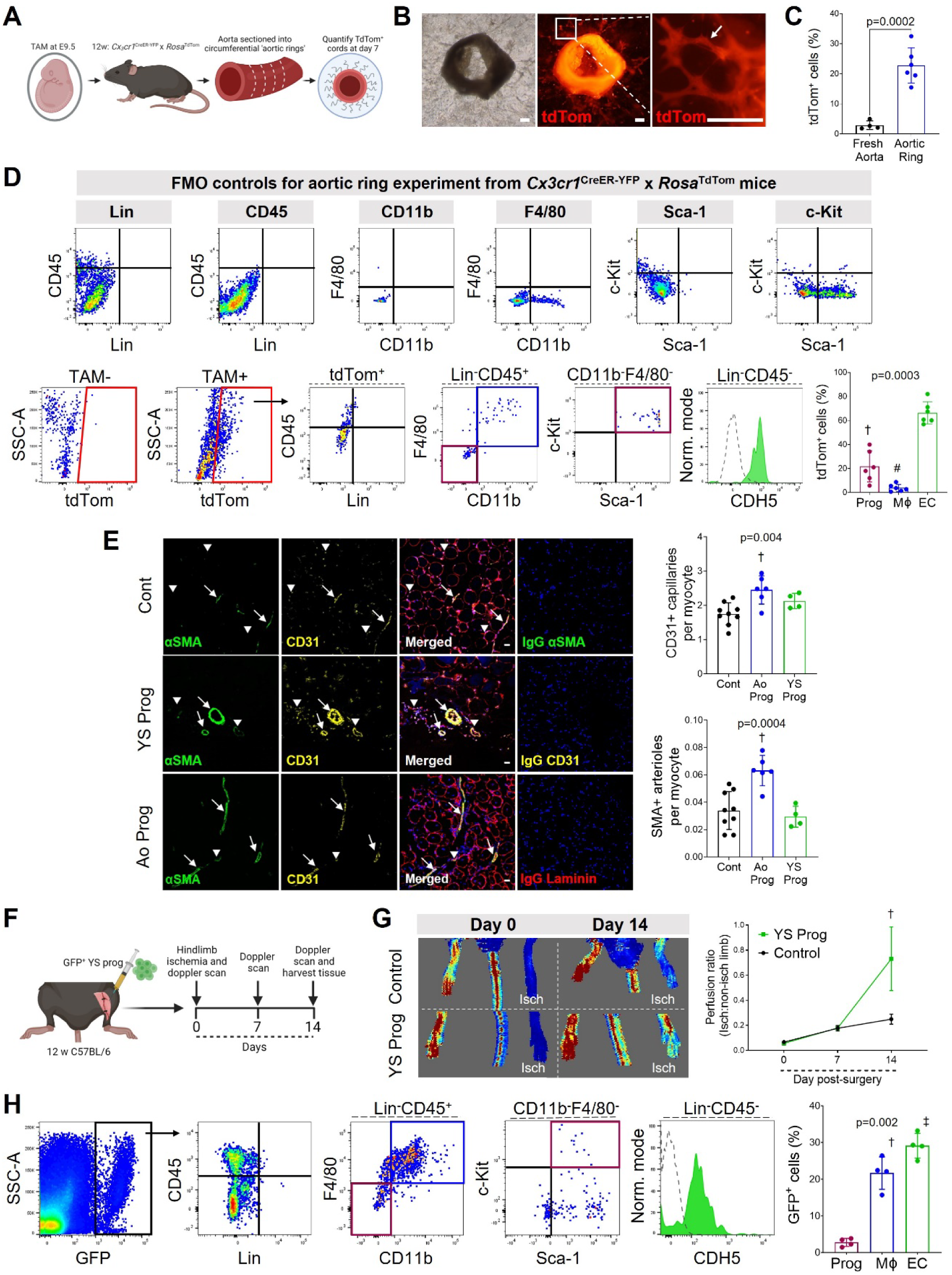
Endothelial and macrophage differentiation capacity of YS-derived progenitors. (A) Schematic for aortic ring assay performed using aortas from E9.5 TAM-induced adult *Cx3cr1*^CreER-YFP^ x *Rosa*^tdTom^ mice to label embryonically derived cells tdTom^+^. (B) Light and fluorescence microscopy images of aortic ring show tdTom^+^ adventitial sprouting. (C) Graph compares % tdTom^+^ expression (not normalized to microglia) in day 7 aortic ring assays with fresh digests of whole aorta. N=4-6; unpaired t-test. (D) Flow cytometry of aortic ring sprouting assay from 12 w *Cx3cr1*^CreER-YFP^ x *Rosa*^tdTom^ mouse with TAM-induction at E9.5. Top row shows FMO controls. Bottom row shows representative sample, with gates used for macrophages (Mϕ, blue), progenitors (Prog, maroon) and endothelial cells (green histogram, EC; dotted histogram, FMO control). Also shown is the tdTom expression for the no TAM (TAM^−^) control used in this experiment. Graph shows the composition of embryonically derived tdTom^+^ cells after aortic ring sprouting. N=6; repeated measures ANOVA. Multiple comparison test: †p<0.01 and #p<0.0001 vs EC. (E) Confocal microscopy of immunolabeled sections of hindlimb muscle at day 14 after ischaemic surgery and receiving injections of Matrigel^TM^ control (top) or E9.5 YS CFU-M progenitors (middle) or Aortic (Ao) CFU-M progenitors (bottom). Staining with antibodies to CD31 (yellow), αSMA (green), laminin (red) for myocytes and DAPI (blue) for nuclei. IgG negative control staining also shown. Graphs show capillary and arteriolar density. N=4-9/ group, three sections per mouse. One-way ANOVA. †p<0.01 vs control. (F) Schematic of 1⁰ transfer of GFP^+^ YS CFU-M progenitors into hindlimb muscle after hindlimb ischemia surgery with two-week follow-up. (G) Laser Doppler perfusion images of mice on days 0 and 14 after ischemia and receiving cell-free control (above) or YS progenitors (below). Graph shows results of follow-up. N=4 for Prog, n=6 for control. Mixed effects two-way ANOVA: p=0.0004 for time; p=0.08 for group; p=0.02 for time x group. Multiple comparison test: †p<0.01 for Prog vs control. (H) Flow cytometry plots and graph show cells produced by GFP^+^ YS progenitors in recipient hindlimb muscle. n=4. Green histogram, sample; dotted, FMO control. Repeated measures ANOVA. Multiple comparison test: †p<0.01 and ‡p<0.001 vs Prog. Data summarized as mean±SD. Scale bar, 100 μm (B, E). Also see **Fig. 5**.

**Extended data Fig. 11.**
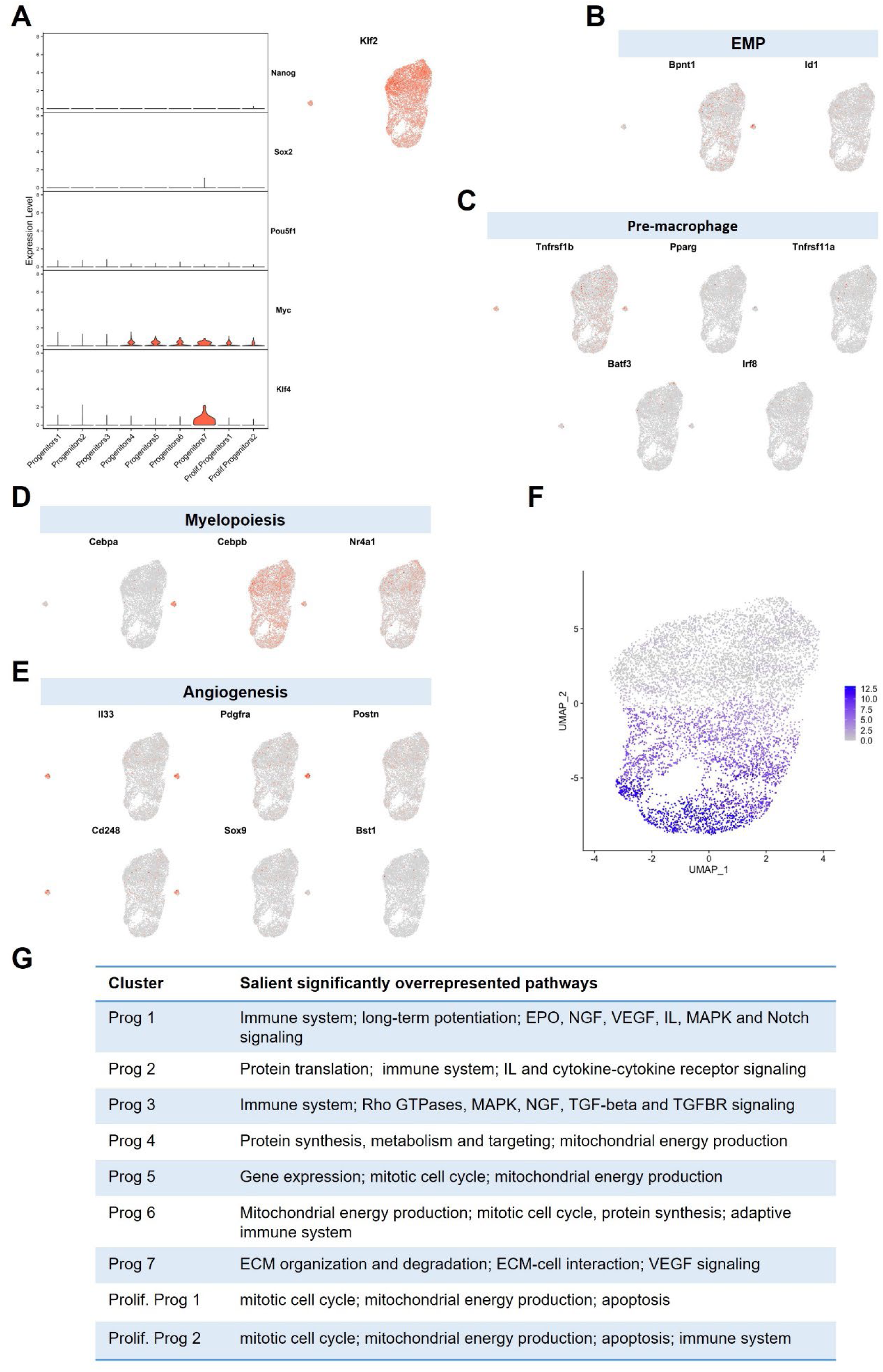
Aortic CFU-M progenitors exhibit a myelopoietic and vasculogenic transcriptional profile with a continuum of cell states. Aortic CFU-M progenitors cultured from two adult C57BL/6J mice were pooled and viable cells analyzed by scRNA-seq. (A) Violin plots illustrate cluster-wise expression of selected stem cell genes (left) and UMAP plot overlaid with expression of *Klf2* (right) showing cell specificity of expression. (B-E) Expression levels of the indicated marker genes for B) EMPs, C) pre-macrophages, D) myelopoiesis and E) angiogenesis overlaid on the UMAP plot showing cell specificity of expression. Expression levels in A-E are shown as log normalised counts. The grey to red gradient represents low to high expression. (F) UMAP plot showing trajectory analysis of single cells in pseudotime. Cells are colored by pseudotime. The analysis showed progression of cell states from cluster 1 at the top to proliferative clusters at the bottom. The Monocle3 algorithm identified Cluster 7 as a separate partition, thus it was not included in the analysis. (G) The table lists the notable overrepresented pathways in the assigned cell clusters. Pathway analysis was performed using InnateDB. EPO, erythropoietin; NGF, nerve growth factor; VEGF, vascular endothelial growth factor; IL, interleukin; MAPK, mitogen activated protein kinase; TGF, transforming growth factor; TGFBR, transforming growth factor-beta receptor; ECM, extracellular matrix. Also see **Fig. 6**.

